# Prenatal Methadone Exposure Leads To Disruptions In Adult-Born Dentate Granule Cell Survival And Female Persistent Fear Responding

**DOI:** 10.1101/2023.06.20.545764

**Authors:** Meredith E. Gamble, Marvin R. Diaz

## Abstract

Methadone is used for the treatment of opioid use disorder, including in pregnant patients. Research has established several consequences of prenatal exposure to misused opioids, however little work has investigated the effects of prenatal methadone exposure (PME) on the offspring long-term, despite the continued prescription to pregnant individuals. The current study aimed to identify the long-term cognitive impairments arising from PME and assess hippocampal neurogenesis in these adult offspring. Pregnant Sprague Dawley rats were injected with methadone or sterile water twice daily from gestational day 3-20 or were left undisturbed as naïve controls. Adult offspring were tested in one of three behavioral tasks to assess pattern separation, spatial learning and memory, and contextual learning and memory, or were assigned to hippocampal tissue collection. For assessment of neurogenesis, offspring underwent injections of bromodeoxyuridine, and brains were collected at 24hr, 2wks, or 4wks for immunofluorescent staining. Methadone-exposed females, but not males, showed subtle impairments in pattern separation and heightened freezing during the extinction period in the fear conditioning task, and spatial memory in both sexes remained unaffected. Additionally, PME did not alter the rate of dentate granule cell proliferation but did significantly reduce the number of adult-born neuron surviving to a mature phenotype in the PME females at the 4wk timepoint. This work adds to the understanding of PME on offspring long-term and demonstrates female-specific sensitivity to these consequences. Future work is needed to fully investigate the neural disruptions arising from PME, with the goal of better supporting exposed individuals long-term.

## 1. Introduction

With the rise in opioid use among pregnant persons and subsequent cognitive difficulties in exposed children, it is crucial to determine the neural and developmental disruptions that may contribute to the trajectory of deficits in these children. The hippocampus is one major regulator of many learning- and memory-associated behaviors, and importantly, the hippocampus is also a target of prenatal opioid exposure (POE)-induced dysfunction throughout the lifetime. As reviewed by Simmons and colleagues (2022), POE has detrimental effects on synaptic plasticity throughout the brain and across development [1]. It is suspected that the hippocampus may be particularly sensitive to deleterious effects of opioid exposures, particularly because synaptic plasticity in this region is regulated by opioid co-release with glutamate for the induction of long-term potentiation [1–4]. This has been established predominantly with preclinical models, showing impairments in synaptic plasticity, aspects of neurogenesis, and reductions in markers of neuronal health and survival [5–13]. These factors are fundamental in maintaining proper brain function and healthy learning and memory maintenance. However, while indirect measurements of neuronal health and survival have been used, such as expression of brain-derived neurotrophic factor (BDNF) [14–17], a more direct measurement of neurogenesis following POE is necessary to continue to fully understand the disrupted functions within the hippocampus.

Additionally, this body of research relating to POE effects on the hippocampus has predominantly focused on exposure to misused opioids, such as morphine and oxycodone, but limited research has examined the impacts of prenatal exposure to commonly prescribed medications for the treatment of opioid use disorder (MOUDs), such as methadone. One recent study that used a prenatal methadone model found significant reduction in both BDNF and GAD67 in the hippocampus of adolescent rats, with females showing greater reductions [18], indicating decreased GABA function and cell health in the hippocampus of these young offspring. Additionally, a study conducted by Wu and colleagues found that prenatal exposure to buprenorphine, an alternative pharmacotherapy to methadone, resulted in decreased neurogenesis in cultured neurospheres from the prefrontal cortex and reduced BDNF in weanling-age rats [11]. This suggests that prenatal exposure to these opioid therapies similarly disrupts typical neural development and neuronal proliferation processes, and therefore, should continue to be studied in these contexts, in light of the continued prescription of MOUDs for use in pregnant persons [19–21].

As discussed, these hippocampal impairments are consistent with the significant cognitive deficits that are prominent in prenatal opioid-exposed children as they develop. And while the hippocampus is not the sole region regulating these behaviors, identification of the ways that POE disrupts this region is essential for continuing to probe the brain circuits and networks involved in producing various learning and memory deficits. Several studies have shown differences in spatial learning and memory using the Morris water maze in offspring following prenatal morphine and oxycodone exposures, including impaired acquisition of spatial learning in adolescent and young adult (∼ postnatal day [P] 51) rodents following prenatal morphine exposure [5, 22]. A recent study also showed impairments in Morris water maze spatial learning only in adult male offspring, with no effect in adult females, following developmental morphine exposure [23]. Importantly, however, several recent studies modeling prenatal methadone or buprenorphine exposures have found no changes in Morris water maze performance in weanling age offspring [24], male offspring with no age listed [25], and impairments in P70 offspring following PME, but not prenatal buprenorphine exposure [26].

This lack of impairments would support the advantages of using these MOUDs during pregnancy compared to continuance of misused opioids, but further assessments are needed to fully understand the nuances of effects for each of these treatments on cognition and which may be preferential for use during pregnancy.

Two other cognitive behavioral tasks that are less commonly studied under POE conditions are contextual fear conditioning and pattern separation. Both of these behaviors model essential cognitive functions, including the ability to inhibit a fear response previously paired with the environment and discrimination of similar stimuli and experiences into distinct representations. Tan and colleagues have shown impaired contextual fear acquisition and extinction in adult male offspring following prenatal morphine exposure; however, this study did not include female offspring [8], so further assessment is required. Furthermore, pattern separation is an essential skill that is commonly linked to impaired adult hippocampal neurogenesis, but to our knowledge, very few studies have examined how POE may disrupt performance on these tasks. One study that used the novel object location task in prenatal heroin-exposed adult offspring showed that while the heroin-exposed offspring explored the novel location object less than saline-exposed, they still preferred the object in the novel location, indicating no major impairments in this task [27]. However, again, only male mice were used in this study, leaving a critical gap in knowledge. Therefore, our study addresses a major gap in knowledge on contextual learning and memory and pattern separation abilities in adult offspring following POE, notably with results from both sexes.

In previous work [28], we have shown altered novel object exploration and impaired working spatial memory in adult female offspring as a result of PME, accompanied by hippocampal dentate granule cell (DGC) dysfunction in female offspring. And while inhibition of hippocampal DGCs in PME females may underlie some of the identified cognitive deficits, the males did not show any significant disruptions to DGC function. This suggests that other hippocampal mechanisms that regulate cognition may be impaired by this exposure. Therefore, in this research, we examined if PME leads to reduced proliferation and integration of DGCs overall. As such, we have assessed the role of PME on the process of adult hippocampal neurogenesis. Using bromodeoxyuridine (BrdU) and immunofluorescence, we have labeled proliferating cells and tracked them through three timepoints using the cellular markers that are expressed throughout the maturation of newborn cells. This has allowed us to track not only the number of new cells that are being produced, but also how many are surviving to a mature phenotype. Additionally, as discussed above, we have further assessed cognitive impairments in adult offspring by adding three behavioral tasks: Novel Object in Place for assessment of pattern separation, Morris water maze for assessment of spatial learning and memory, and contextual fear conditioning for assessment of context learning and memory. With this information, we add new knowledge on the long-term effects of PME within the hippocampus, as well as a more thorough understanding of the long-term cognitive impairments that may arise as a result of this exposure.

## 2. Methods

### 2.1 Animals

Male and female Sprague-Dawley rats were bred in-house, with breeding pairs originating from Envigo (Indianapolis, IN). Following exposure procedures, data from a total of 297 (145 male and 152 female) offspring from 41 litters (13 naïve, 13 water, and 15 methadone) were used (see **Supplementary Table 1** for breakdown of number of animals, litters, and animals per litter for each experiment). At weaning (P21), same-sex littermates were housed two to three per cage and randomly assigned to an experimental condition. Rats continued to receive food (5L0D PicoLab Laboratory Rodent Diet) and water *ad libitum* and were maintained on a 12-hr light-dark cycle (lights on at 07:00). All rats were treated following guidelines for animal care under protocols approved by Binghamton University Institutional Animal Care and use Committee.

### 2.2 Breeding and Prenatal Methadone Exposure (PME)

Breeding and exposure procedures were identical to those described previously [28]. To summarize, two naïve adult females were paired with one naïve adult male for up to four days. Pregnancy was confirmed with vaginal smears, and pregnant females were single-housed and randomly assigned to an exposure condition: naïve control, water-injected control, or methadone. PME began on G3, with dams receiving twice daily subcutaneous injections, 12 hours apart, of 5 mg/kg (G3) and 7 mg/kg (G4-20) of methadone dissolved in sterile water or sterile water alone. Methadone- and water-exposed dams were moved to cages with no bedding for three hours after each injection for monitoring and prevention of ingesting bedding material. After three hours of monitoring, dams were returned to homecage until the next injection. Naïve dams were left undisturbed in their homecage throughout gestation except for daily weights. All dams were left undisturbed after G20 to give birth. Pups were weighed, sexed, and culled to 12 (equal sex ratio where possible) on P2, weighed on P7 and 12, and weaned into cages with same-sex littermates on P21. Following weaning, offspring were left undisturbed until adult testing.

### 2.3 Novel Object in Place

Following testing in a Spontaneous Alternation task (results reported in previous work [28]), the same animals (n = 8 per group, per sex) were tested using the Novel Object in Place (NOP) task. Animals were put through both tasks because Spontaneous Alternation is an exploratory task, with no training conducted, so it is not expected that testing would impact subsequent NOP behavioral testing. These were the only behavioral experiments in which animals were reused.

Testing consisted of two habituation days, two familiarization days, and one test day. For habituation, animals were placed individually in an empty black plexiglass arena (38.5 × 70 × 36 cm), with no objects present, and allowed to freely explore for 10 minutes. After 10 minutes, they were returned to their homecage. Habituation procedures were repeated 24 hours later (Day 2). On Day 3, 24 hours after the second habituation day, animals were again placed in the same plexiglass arena for familiarization days and allowed to freely explore two identical wooden blocks (6.5 x 6.5 x 2.5 cm; 15.28 cm from the furthest wall and 20.9 cm apart) for 6 minutes.

Familiarization day procedures were repeated on Day 4. Both familiarization sessions were recorded and scored to ensure no side preference and adequate exploration of the objects for encoding of the location. Finally, on Day 5 (Test Day), animals were again placed in the same arena and in the same starting position facing the objects, with the left object now moved forward to a new location (32.25 cm from the closest wall, still equidistant to the object in the original location; **Supplementary Fig. 1A**). Animals were allowed to explore the arena and objects for 6 minutes. The test day was recorded and scored for time spent exploring the objects and a discrimination ratio was calculated (time spent investigating the object in the new location divided by the total time investigating both objects on test day). Increased time exploring the object in the new location was used as an indicator of better pattern separation abilities.

### 2.4 Morris Water Maze

A separate cohort of adult offspring (26 males [8 naïve, 8 water, 10 PME] and 28 females [10 naïve, 8 water, 10 PME]) underwent assessment of spatial learning and memory through performance in the Morris water maze with surrounding spatial cues using procedures adapted from previous guidelines [29, 30]. This behavioral assay was selected because it is considered the gold standard for spatial memory assessment in rodent models. A round, green pool (diameter = 182.88 cm, depth = 63.5 cm) was filled with water (22 ± 1° C) to a depth of 33 cm, with a green platform just below the water surface that remained in the same place in the NW quadrant throughout training (**Supplementary Fig 1B**). The water was colored with green paint, so the platform could not be seen. A variety of spatial cues (black paper shapes) were placed on surrounding walls, and the experimenter remained in the same location in the room for all training and test days. Animals were handled for two days prior to the first training session. For training sessions, each animal was brought into the testing room, starting at approximately 15:00 each day, and placed in the water using alternating starting positions for each trial (**Supplementary Fig 1B**). Each training session consisted of 4 trials, for a total of 6 training days, 24 hr apart. Each trial lasted until the animal either located the platform or up to one minute; if the platform was not located in one minute, the experimenter guided the animal to swim to the platform. Once on the platform, each animal was given a 30 s intertrial interval before being removed from the platform and starting the next trial. At the end of 4 trials, animals were dried off and returned to their homecage until the next day. Following the 6 training days, animals went through 2 test days: one 24 hr after the last training session and one 7 days after the last training session. Test days consisted of a one-minute probe trial with the platform removed.

All training and test trials were recorded with an overhead camera and stored for later analyses. Videos were uploaded with experimenter blind to animal conditions and AnyMaze (Stoelting Co., Wood Dale, IL) was used to score acquisition on training days (latency to find the platform, average speed, path length, thigmotaxis, and percent of time spent in the NW quadrant) and spatial memory in the probe tests (number of platform location crossovers, latency to first platform location crossover, average speed, and percent of time spent in the NW quadrant).

During training days, decreased latency and path length to find the platform and higher percent of time in the NW quadrant are indicative of faster acquisition. During the probe tests, a higher number of platform location crossovers, decreased latency to first platform crossover, and increased time spent in the NW quadrant during the probe tests are indicative of better spatial memory.

### 2.5 Contextual Fear Conditioning

Contextual fear conditioning was used to assess context learning in a separate cohort of adult offspring (n = 8 per group, per sex). Following three days of handling, animals were placed in individual fear conditioning chambers (32 × 25 × 25 cm, MED-VFC-SCT-R; Med Associates Inc, Fairfax, VT, USA), made of clear polycarbonate, white acrylic, and stainless-steel floors (shock grids made up of 19 parallel rods, 1 cm apart, with a drop pan below) in sound attenuating cabinets. Chambers were lit with a 20-watt light bulb and were recorded using a near-infrared camera. Chambers were run four at a time, starting at approximately 08:30 each day. On Day 1, animals were left for a 5-minute acclimation period, then received 3 shocks (0.8 mA, 1 s each, 90 s apart) and then were left for an additional 5-minute encoding period before being returned to their home cage. Twenty-four hours later, animals were placed in the same boxes again for 20 minutes, with no shocks administered, to test for retention of conditioned fear as indicated by percent freezing. This test day procedure was repeated three more days (24 hr apart) for assessment of extinction learning of conditioned fear (**Supplementary Fig. 1C)**. The same order of animals was used each day so that each animal was placed in their same respective box every day. Each box was cleaned between each animal using tap water and paper towels. After the final extinction session, chambers were thoroughly cleaned with ethanol. Percent freezing was scored by Med Associates freezing software, with freezing behavior defined as no movement for at least 18 of 30 frames per second.

### 2.6 Adult Neurogenesis Experimental Design

A total of 74 males (25 naïve, 25 water, 24 PME) and 78 females (26 naïve, 28 water, 24 PME) underwent injections for labeling of cell proliferation and brain collection for immunofluorescent staining. Brains were collected at 3 different timepoints to assess cell proliferation and survival: 24 hr after the final injection (co-labeling with glial fibrillary acidic protein [GFAP]) for general proliferation, 2 weeks after injection day (co-labeling with doublecortin [DCX]) for assessment of immature phenotype of adult-born neurons, and 4 weeks after injection day (co-labeling with neuronal nuclei [NeuN]) for assessment of a mature phenotype of adult-born neurons **(Fig. 1)**.

**Figure 1.**
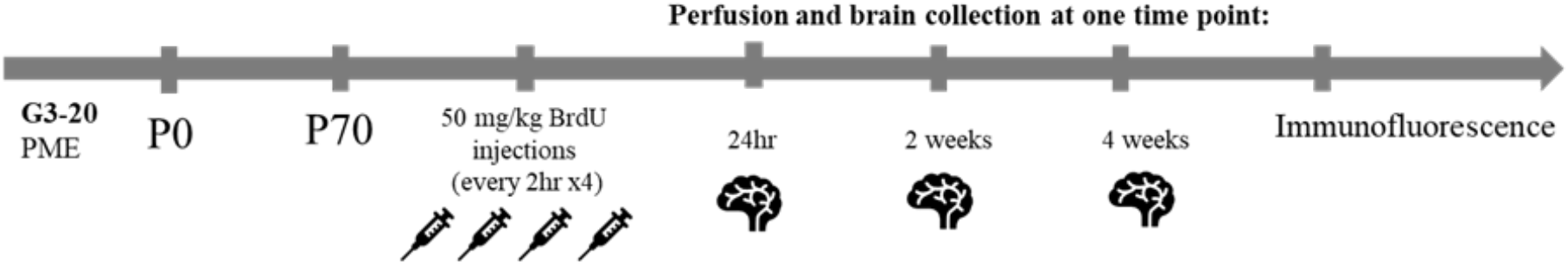
Timeline of BrdU experiments. Following postnatal day 70, male and female offspring received four 50mg/kg BrdU injections, one every 2 hrs, and were perfused at one of three timepoints (24hrs, 2 wks, or 4wks) for tissue collection and immunofluorescent processing.

### 2.7 Bromodeoxyuridine (BrdU) Injections and Tissue Collection

All animals received four 50 mg/kg intraperitoneal injections, 2 hr apart, of BrdU (Sigma CAS No: 59-14-3; Product no. 10280879001). BrdU solution (15 mg/mL BrdU in sterile saline and 1% DMSO) was prepared fresh each morning, and saline was heated to 37°C. Once in solution, BrdU remained at room temperature for the rest of the day and was disposed of using proper hazardous waste handling. All animals were weighed in the morning before the first injection. After each injection animals were returned to their homecage, and experimenters monitored daily weights and general care for up to 72 hrs following injections. For the 2- and 4- week groups, cages were changed after 72 hrs in full personal protective equipment out of an abundance of caution for any excreted BrdU in the cage. At this point, animals were left undisturbed in homecages with normal cage changes until their brain collection timepoint.

At each timepoint, animals were anesthetized with pentobarbital and transcardially perfused with ice cold phosphate-buffered saline (PBS) followed by 4% paraformaldehyde (PFA). Brains were extracted and post-fixed overnight (16-20 hours) in 4% PFA at 4°C. They were then transferred to a 30% sucrose in PBS solution at 4°C for approximately one week before flash freezing at -20 °C in methyl butane and stored at -20 °C until sectioning. 40 μm slices were made of the entire dorsal hippocampus (dHPC) at -20 °C using a cryostat (CM1510 Leica Biosystems). Sections were stored individually in 96-well plates containing antifreeze solution (30% ethylene glycol, 30% glycerol in PBS solution) at -20 °C until immunofluorescent staining.

### 2.8 Immunofluorescent Staining and Imaging

For immunofluorescent staining, every sixth slice from the entire dHPC was selected, for a total of six slices per animal. Slices were washed three times in PBS, immediately followed by pretreatment for BrdU denaturing in 2N hydrochloric acid for 15 min at 37°C. Slices were then neutralized in borate buffer (pH: 8.62) at room temperature for 10 min, followed by another three, 5-minute PBS washes. Following this pretreatment, slices were incubated in blocking buffer (5% normal donkey serum, 0.1% Triton-X in PBS) for one hour at room temperature on an orbital shaker. After blocking, slices were incubated in primary antibody cocktail overnight (∼16 hr) on an orbital shaker at 4°C. Primary antibody cocktails were made in blocking buffer and consisted of mouse anti-BrdU primary antibody and one of the following (respective of timepoint): rabbit anti-GFAP, rabbit anti-DCX, or rabbit anti-NeuN (**Table 1**). All immunofluorescence experiments were sufficiently piloted with control experiments (primary omissions, BrdU^-^ tissue, and optimal dilutions) to ensure accurate labeling of target markers before beginning with experimental tissue.

**Table 1.** Immunofluorescent Antibodies. Immunofluorescent antibodies, dilutions used, and supplier information. BrdU = bromodeoxyuridine, GFAP = glial fibrillary acidic protein, DCX = doublecortin, NeuN = neuronal nuclei, DAPI = 4’,6-diamidino-2-phenylindole

| **Antibody** | **Dilution** | **Supplier and Purchase Number** |
| --- | --- | --- |
| Mouse anti-BrDU Monoclonal Antibody | 1:250 | Invitrogen, MA3-071 |
| Rabbit anti-GFAP Polyclonal Antibody | 1:1000 | Dako, Z033401-2 |
| Rabbit anti-DCX Polyclonal Antibody | 1:800 | Cell Signaling Tech, 4604 |
| Rabbit anti-NeuN Polyclonal Antibody | 1:1000 | Invitrogen, PA5-78499 |
| Donkey anti-Mouse IgG Secondary Antibody (Alexa Fluor 594) | 1:500 | Invitrogen, A-21203 |
| Donkey anti-Rabbit IgG Secondary Antibody (Alexa Fluor 680) | 1:500 (GFAP, NeuN)<br>1:1000 (DCX) | Jackson ImmunoResearch<br>711-625-152 |
| ProLong Gold Antifade Mountant with DAPI | - | Invitrogen, P36931 |

**Table 1.** Immunofluorescent antibodies, dilutions used, and supplier information. BrdU = bromodeoxyuridine, GFAP = glial fibrillary acidic protein, DCX = doublecortin, NeuN = neuronal nuclei, DAPI = 4',6-diamidino-2-phenylindole
| Antibody | Dilution | Supplier and Purchase Number |
| --- | --- | --- |
| Mouse anti-BrdU Monoclonal Antibody | 1:250 | Invitrogen, MA3-071 |
| Rabbit anti-GFAP Polyclonal Antibody | 1:1000 | Dako, Z033401-2 |
| Rabbit anti-DCX Polyclonal Antibody | 1:800 | Cell Signaling Tech, 4604 |
| Rabbit anti-NeuN Polyclonal Antibody | 1:1000 | Invitrogen, PA5-78499 |
| Donkey anti-Mouse IgG Secondary Antibody (Alexa Fluor 594) | 1:500 | Invitrogen, A-21203 |
| Donkey anti-Rabbit IgG Secondary Antibody (Alexa Fluor 680) | 1:500 (GFAP, NeuN)<br>1:1000 (DCX) | Jackson ImmunoResearch<br>711-625-152 |
| ProLong Gold Antifade Mountant with DAPI | - | Invitrogen, P36931 |

Following primary incubation, slices went through three, 5-minute washes in PBS and incubated in secondary antibody cocktail for one hour at room temperature on a shaker.

Secondary antibody cocktails were made in blocking buffer and contained donkey anti-mouse (BrdU reporter) and donkey anti-rabbit (GFAP, DCX, or NeuN reporter) **(Table 1).** Following incubation, slices were washed in PBS and mounted on charged slides using ProLong Gold with DAPI mounting media. Slides were left flat to dry overnight at room temperature, coverslips were then sealed with clear nail polish, and slides were stored in a dark slide box at 4°C until imaging (approximately one week later). Slides were imaged using a slide scanning microscope (Olympus Slideview VS200) at 20X captured with Extended Focal Imaging (EFI).

### 2.9 Cell Counting

Unbiased cell counts were completed using HALO software (Indica Labs) with the experimenter blind to experimental conditions. Cell identification was accomplished by creating a cell recognition protocol specific for the experiment to distinguish fluorescent signal from noise. For each EFI image, individual cells were identified with a DAPI^+^ phenotype. The protocol then quantified the presence of each fluorophore within the DAPI^+^ region with a set 0.3 μm cytoplasm radius around it. Output included total cell counts and percent of each phenotype (BrdU^+^ alone or with colocalization with secondary marker [GFAP, DCX or NeuN]). Cell counts were completed for both hemispheres of each slice for all six slices per animal. Analysis was restricted to the dentate gyrus of the dHPC in accordance with Paxinos and Watson [31] rodent atlas, with an ∼100 μm total width around each blade. In cases where both hemispheres did not make it through processing, cell counts from the available hemisphere were doubled. In cases where only five slices for the animal made it through processing, the sixth slice value was replaced with the average of the other five slices for that animal. Total cell counts were summed for a total value for each animal for each phenotype (BrdU^+^ total, BrdU^+^ colabeled with the secondary marker [BrdU^+^/GFAP^+^ at 24hr, BrdU^+^/DCX^+^ at 2 weeks, or BrdU^+^/NeuN+ at 4 weeks], and the totals for each secondary marker independently [GFAP^+^, DCX^+^, NeuN^+^, respectively]). Since these values represent the totals from every sixth slice, with a total of six slices, they can be used as an estimate of 1/6^th^ the total dHPC dentate gyrus values.

### 2.10 Data Analyses

We have analyzed each sex independently for all experiments in this study. This is due to the substantial literature on sex differences in spatial navigation, pattern separation, and contextual fear conditioning tasks [32–38], as well as in adult hippocampal neurogenesis processes [38–41], and hippocampal dentate granule cell electrophysiological processes [28].

Statistical analyses for all data were completed using Prism 8 (GraphPad, San Diego, CA). Rounded p-values ≤ 0.05 were considered statistically significant. All data were tested for normality (Kolmogrov-Smirnov test), and nonparametric tests were used when data were not normally distributed. NOP data were analyzed using a one-sample t-test comparing mean discrimination ratio values to 0.5 (chance), and data were followed-up with a one-way ANOVA to test for general group differences in discrimination ratios. NOP data were also analyzed by cumulative minutes to identify if exploration behavior only occurred during initial few minutes, but this did not change results, so data are reported for the full 6 minutes. Female data did not pass the normality test, so data were analyzed using Wilcoxon signed-rank test. Morris water maze data from training days were analyzed using a repeated-measures two-way ANOVA (exposure x day) for each parameter with Geisser-Greenhouse correction for sphericity where applicable; each retention test day (24 hr and 7 day) were analyzed separately with one-way ANOVAs for each parameter. Fear conditioning data were analyzed using repeated-measures two-way ANOVAs, with the shock day and test day each analyzed separately across minute (exposure x minute), as well as the test day and three extinction days analyzed together across day (exposure x day), since the testing and extinction days were all procedurally the same.

Immunofluorescence data were analyzed with a one-way ANOVA for each parameter, with data from each of the three timepoints analyzed separately. For sex comparisons in naïve groups, unpaired t-tests were used comparing male and female data for each timepoint. Tukey’s post hoc with correction for multiple comparisons was used for follow-up on any significant effects.

## 3. Results

### 3.1 PME does not impair pattern separation abilities in adult offspring

Adult offspring underwent five days of testing using the NOP task for assessment of pattern separation. Each animal completed 2 days of habituation, 2 days of familiarization with the objects, and 1 test day, as described above. Investigation of the objects was recorded and scored for both familiarization days to ensure adequate investigation of objects and no side preference before the test day. Neither males nor females showed any significant differences across exposure groups for time spent interacting with the objects during the familiarization days (females: *F*(2,21) = 0.209, *p* = 0.813; males: *F*(2,21) = 0.619, *p* = 0.548) and no significant difference in discrimination ratios during these training days (females: *F*(2,21) = 0.831, *p* = 0.449; males: *F*(2,21) = 0.784, *p* = 0.469) (**Supplementary Fig. 2).**

For the test day, discrimination ratio data from methadone females were not normally distributed, so a Wilcoxon Signed Rank Test was used. Naïve females showed a preference for the object in the novel location significantly greater than chance (theoretical mean = 0.5) (*Mdn =* 0.553, *p* = 0.023), while water- and methadone-exposed offspring did not differ from chance (water: *Mdn =* 0.572, *p* = 0.078; methadone: *Mdn =* 0.383, *p* = 0.312). Female offspring showed a trend toward significant differences in discrimination ratios across exposure groups (*H*(2) = 5.420*, p* = 0.067) **(Fig. 2A).**

**Figure 2.**
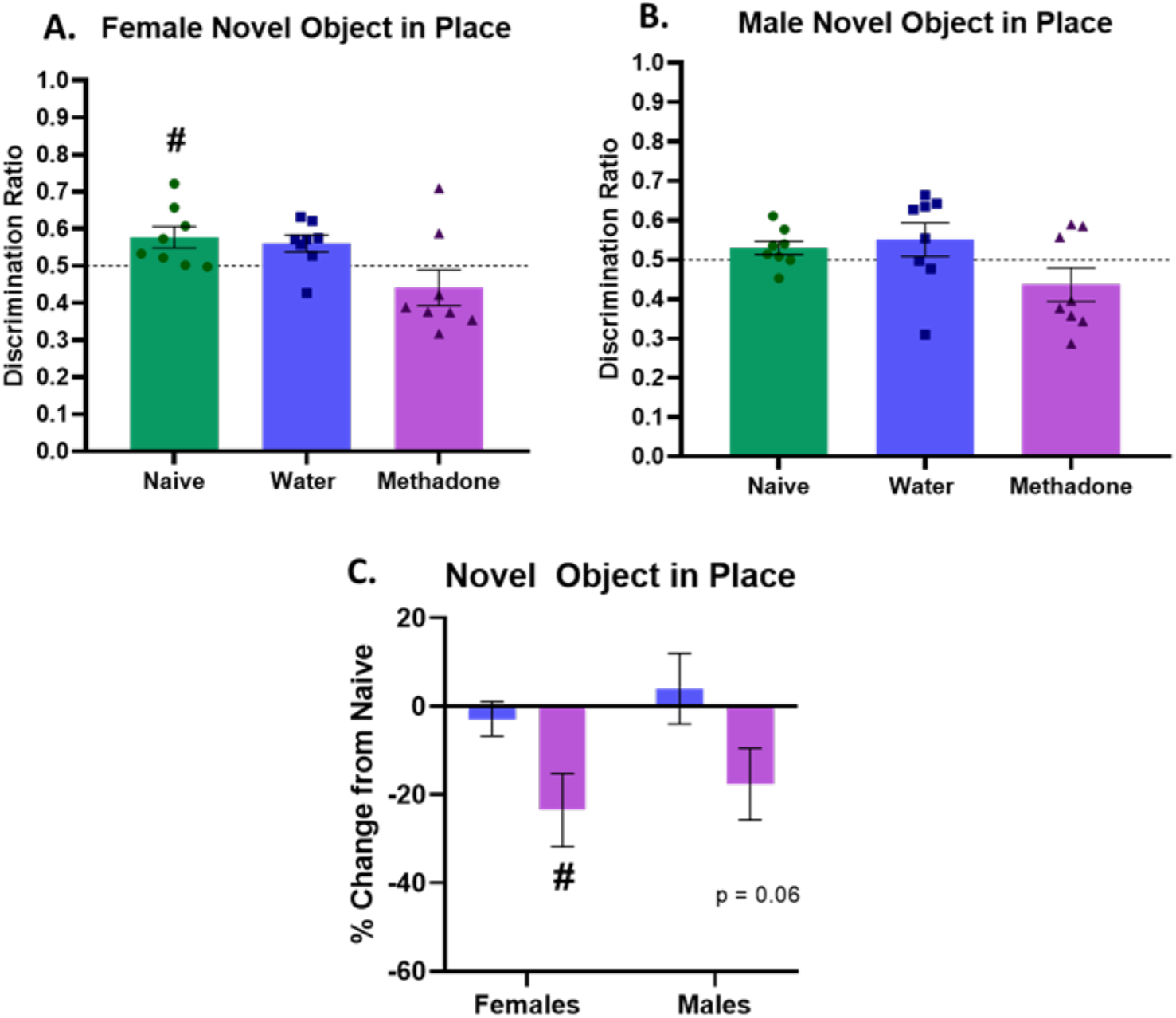
PME does not significantly impair pattern separation abilities. In females, naïve offspring investigated the object in the novel location significantly more on test day, whereas methadone- and water-exposed females did not differ from chance (A). No exposure groups in male offspring significantly differed from chance (B). As percent change from naïve, PME females showed a significant reduction in investigation of the object in the novel location, and methadone- exposed males showed a strong trend toward the same, with no differences in water-exposed offspring of either sex. Data points represent values from each animal; error bars represent SEM. # = *p* < 0.05 compared to theoretical mean (discrimination ratio = 0.5, percent change = 0).

For male offspring, no exposure groups had discrimination ratios that significantly differed from chance (theoretical mean = 0.5) (naïve: *t*(7) = 1.732, *M* = 0.530, *p* = 0.127; water: *t*(7) = 1.203, *M* = 0.551, *p* = 0.268; methadone: *t*(7) = 1.486, *M* = 0.436, *p* = 0.181).

Additionally, males did not show any significant differences in discrimination ratios across the three exposure groups (*F*(2,21) = 2.842, *p* = 0.081) **(Fig. 2B).**

Discrimination ratio data have also been shown as percent change compared to naïve controls (**Fig. 2C**). PME females showed significant decreases compared to naïve females (theoretical mean = 0) (*t*(7) = 2.835, *M* = -23.49, p = 0.025), and PME males showed a trend toward significant reductions compared to naïve males (*t*(7) = 2.180, *M* = -17.61, *p* = 0.066), while water-exposed offspring of either sex did not significantly differ compared to naïve offspring (females: *t*(7) = 0.738, *M* = -2.868, *p* = 0.485; males: *t*(7) = 0.501, *M* = 4.006, *p* = 0.632).

### 3.2 PME does not alter spatial learning or memory in the Morris water maze task

Adult offspring were assessed for spatial learning and memory using the Morris water maze task. Each animal underwent 4 training trials per day for 6 days, followed by a 24hr and 7- day retention test. Across the 6 training days, female offspring showed no significant differences across exposure groups for any of the following measures: latency to find the platform (*F*(2,25) = 0.537, *p* = 0.591; **Fig. 3A**), path length to find the platform (*F*(2,25) = 0.639, *p* = 0.537; **Fig. 3B**), or percent of time spent in the platform quadrant (*F*(2,25) = 1.766, *p* = 0.198; **Fig. 3C**). This indicated no significant PME impairments in spatial learning. Animals did show significant differences across days, with all animals making improvements from the first to the sixth training day (latency: *F*(3.368, 84.19) *=* 45.14, *p* < 0.0001; path length: *F*(3.373, 84.32) = 57.08, *p* <

**Figure 3.**
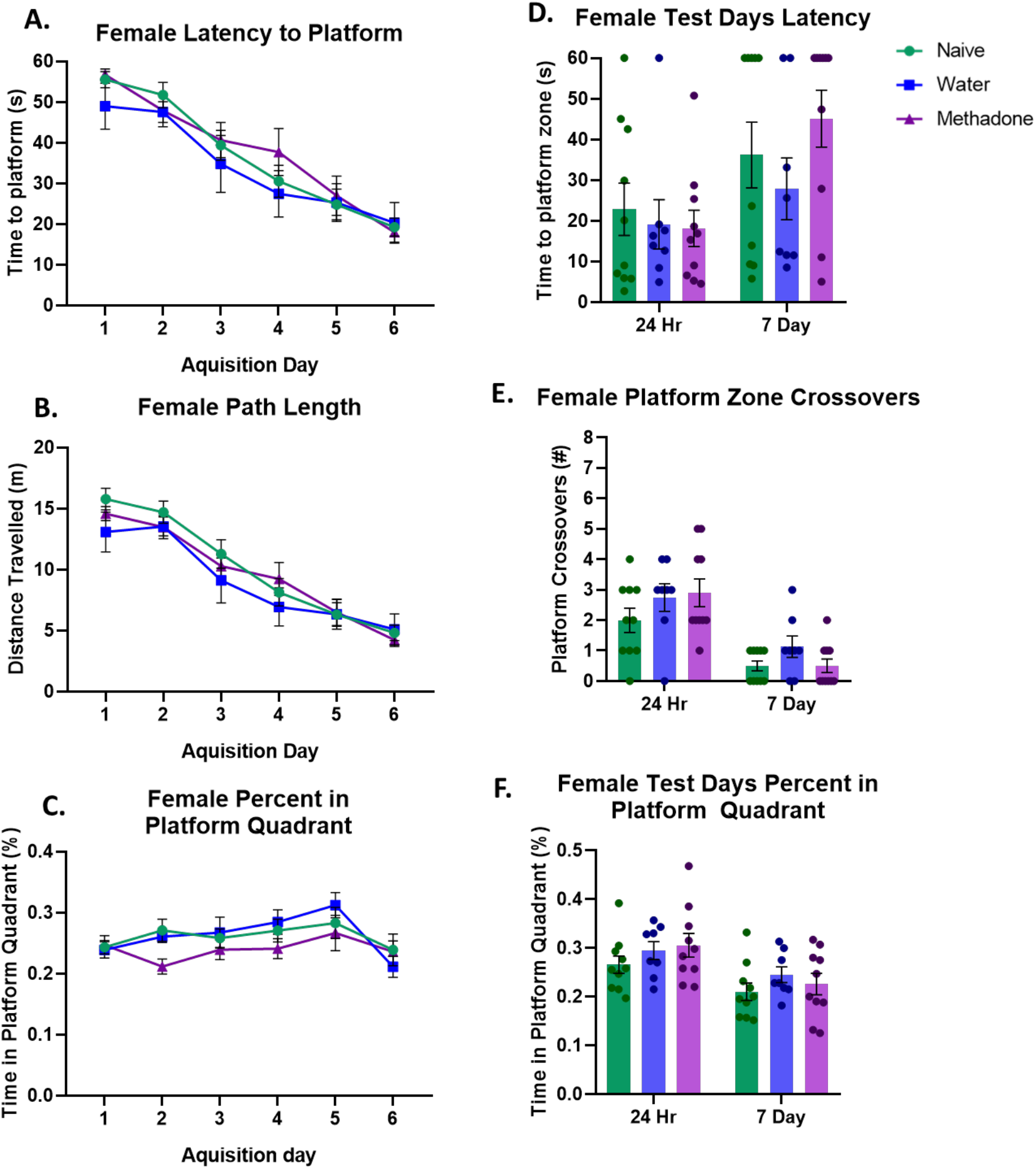
PME does not significantly impair female spatial learning or memory. Female offspring showed no significant impairments in spatial acquisition, measured at time to find the platform (A), path length to the platform (B), and percent of time spent in the platform quadrant (C) across 6 training days. Females also showed no significant impairment in spatial memory, measured as time to the platform zone (D), platform zone crossovers (E), and percent of time in the platform quadrant (F) during the 24hr and 7-day retention tests. Data points represent values from each animal; error bars represent SEM.

0.0001; percent of time in platform quadrant: *F*(3.497, 87.43) = 3.646, *p* = 0.012). There were no significant interactions in any of these measures (latency: *p* = 0.781; path length: *p* = 0.722; percent time in platform quadrant: *p* = 0.512).

For the two test days, females also showed no significant differences in retention across the exposure groups. At the 24hr timepoint, females showed no difference in latency to the first platform zone crossover (*F*(2,25) = 0.198, *p* = 0.822), number of platform zone crossovers in the one minute test period (*F*(2,25) = 1.285, *p* = 0.294) or percent of time spent in the platform quadrant (*F*(2,25) = 1.026, *p* = 0.373). At the 7 day time point, there were also no significant differences in these measures: latency to the first platform zone crossover (*F*(2,25) = 1.238, *p* = 0.307), number of platform zone crossovers in the one minute test period (*F*(2,25) = 2.011, *p* = 0.155) or percent of time spent in the platform quadrant (*F*(2,25) = 0.780, *p* = 0.469) **(Fig. 3D- F)**.

Additional measures included average swimming speed and percent of time engaging in thigmotaxic behavior, as described above. Females showed no significant differences in average speed across the training and test days (exposure: *F*(2,25) = 0.729, *p* = 0.468; interaction: *F*(14,175) = 1.012, *p* = 0.444) and no significant differences in percent of time engaging in thigmotaxic behavior (exposure: *F*(2,25) = 1.270, *p* = 0.298; interaction: *F*(14,175) = 0.379, *p* = 0.979) **(Supplementary Fig. 3).**

For males, offspring showed no significant differences across exposure groups for latency to find the platform (*F*(2,23) = 0.631, *p* = 0.541; **Fig. 4A**) or path length to find the platform (*F*(2,23) = 1.023, *p* = 0.375; **Fig. 4B**). Interestingly, males did show a significant exposure effect in percent of time spent in the platform quadrant (*F*(2,23) = 3.767, *p* = 0.038; **Fig. 4C**). Follow up with Tukey’s post hoc analysis revealed this was driven by the naïve group spending significantly more time in this quadrant compared to both the water-exposed (*p* = 0.014) and methadone-exposed (*p* = 0.030), with no significant difference between methadone and water groups (*p* = 0.870). Animals did show significant differences across days, with all animals making improvements from the first to the sixth training day for latency (*F*(3.865, 88.89) *=* 103.8, *p* < 0.0001) and path length (*F*(4.008, 92.19) = 120.8, *p* < 0.0001), but not percent of time in platform quadrant (*F*(3.880, 89.23) = 1.982, *p* = 0.106). There were no significant interactions in any of these measures (latency: *p* = 0.128; path length: *p* = 0.110; percent time in platform quadrant: *p* = 0.690).

**Figure 4.**
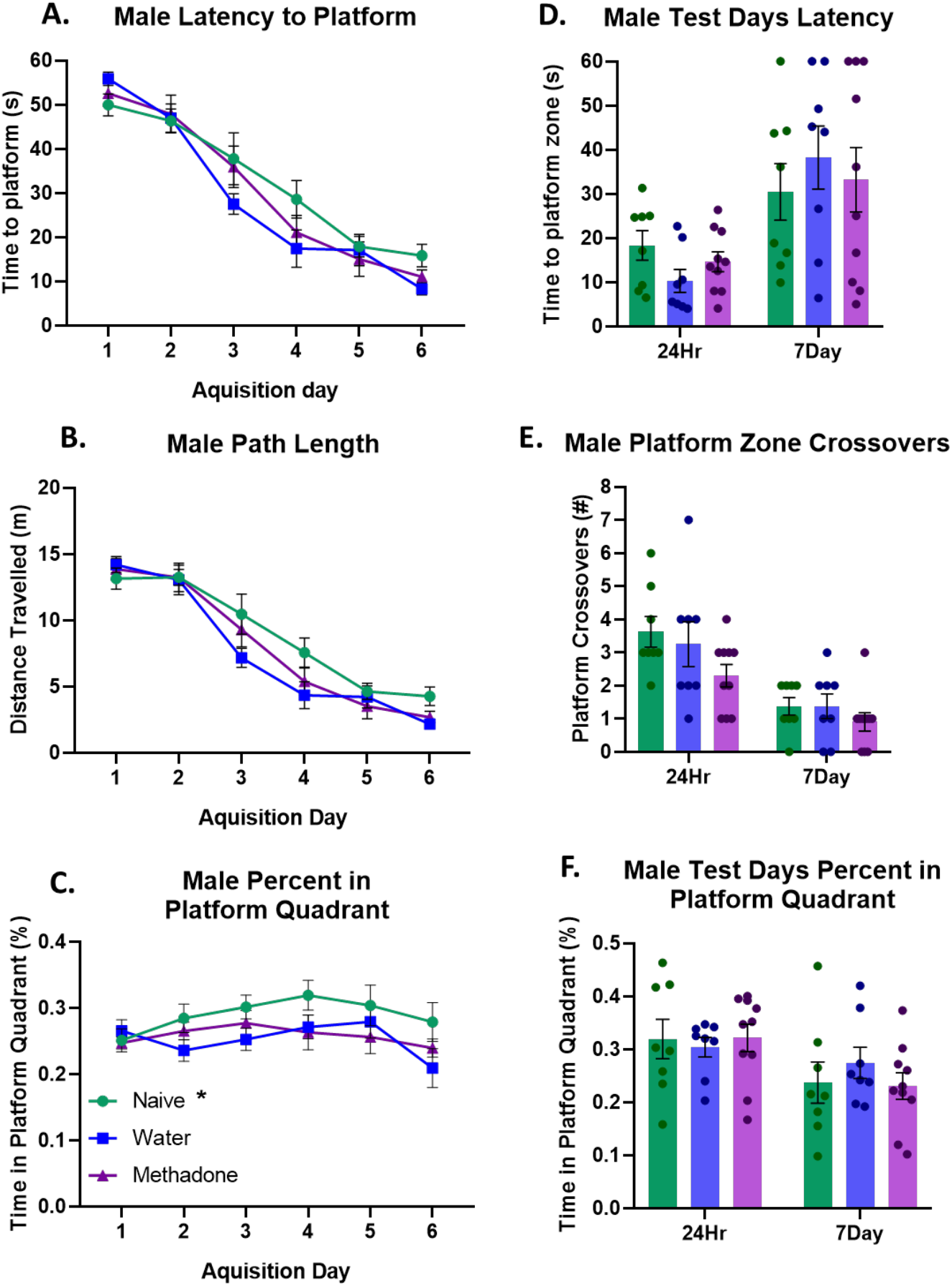
PME does not significantly impair male spatial learning or memory. Male offspring showed no significant differences in time to find the platform (A) or path length to the platform (B) across the 6 training days; however, naïve males did spend significantly more time in the platform quadrant during training (C). Males did not show any significant impairment in spatial memory, measured as time to the platform zone (D), platform zone crossovers (E), and percent of time in the platform quadrant (F) during the 24hr and 7-day retention tests. Data points represent values from individual animals; error bars represent SEM. * p < 0.05 compared to water- and methadone-exposed.

For the two test days, males also showed no significant differences in retention across the exposure groups. At the 24hr timepoint, males showed no difference in latency to the first platform zone crossover (*F*(2,23) = 2.063, *p* = 0.150), number of platform zone crossovers in the one minute test period (*F*(2,23) = 2.065, *p* = 0.150) or percent of time spent in the platform quadrant (*F*(2,23) = 0.113, *p* = 0.894). At the 7 day time point, there were also no significant differences in these measures: latency to the first platform zone crossover (*F*(2,23) = 0.290, *p* = 0.751), number of platform zone crossovers in the one minute test period (*F*(2,23) = 0.856, *p* = 0.438) or percent of time spent in the platform quadrant (*F*(2,23) = 0.570, *p* = 0.573) **(Fig. 4D- F)**.

Males showed a significant interaction in average speed (*F*(14,161) = 2.184, *p* = 0.010), with post hoc results showing this was driven by increased speed in naive controls compared to water males on day 4 (*p* = 0.009) and increased speed in naïve controls compared to methadone on day 5 (*p* = 0.031). There was no significant exposure main effect for average swimming speed across all days (*F*(2,23) = 2.943, *p* = 0.073). Since we did not observe any significant effects on latency to platform and the differences in swimming speed were not significant across all days, we do not believe this explains the differences in the percent of time in platform quadrant measure **(Supplementary Fig. 3).** Finally, males did not show any significant exposure differences in thigmotaxic behavior (exposure: *F*(2.23) = 1.199, *p* = 0.320). However, there was a significant interaction (*F*(14.161) = 2.063, *p* = 0.016), with post hoc tests showing this was driven by an increase in percent of time engaging in thigmotaxic behavior in the naïve group compared to methadone on day 5 (*p* = 0.026). **(Supplementary Fig. 3).**

### 3.3 PME induces persistent heightened freezing in adult female offspring

Adult animals underwent five days of contextual fear conditioning to test contextual fear memory and extinction learning. Each animal went through 1 initial shock day, followed by 1 retention test day and 3 days of extinction learning.

On the shock day, females did not show any increases in percent freezing during the initial acclimation period (5min) and showed substantial increases in freezing once the foot shocks began and during the consolidation period (5min), with a significant main effect of time (*F*(3.701, 77.73) = 222.9, *p* < 0.0001). However, there was no significant exposure main effect in percent freezing across the total 13min on this initial conditioning day (*F*(2,21) = 2.272, *p* = 0.128) or interaction (*F*(24, 252) = 0.817, *p* = 0.715) **(Fig. 5A).** For the test day, animals still showed a significant effect of time, with decreases in percent freezing across the 20min testing period (*F*(6.066, 127.4) = 5.948, *p* < 0.0001), but no significant differences across exposure groups (*F*(2,21) = 1.564, *p* = 0.233), and no significant interaction (*F*(38, 399) = 1.345, *p* = 0.089) **(Fig. 5B).**

**Figure 5.**
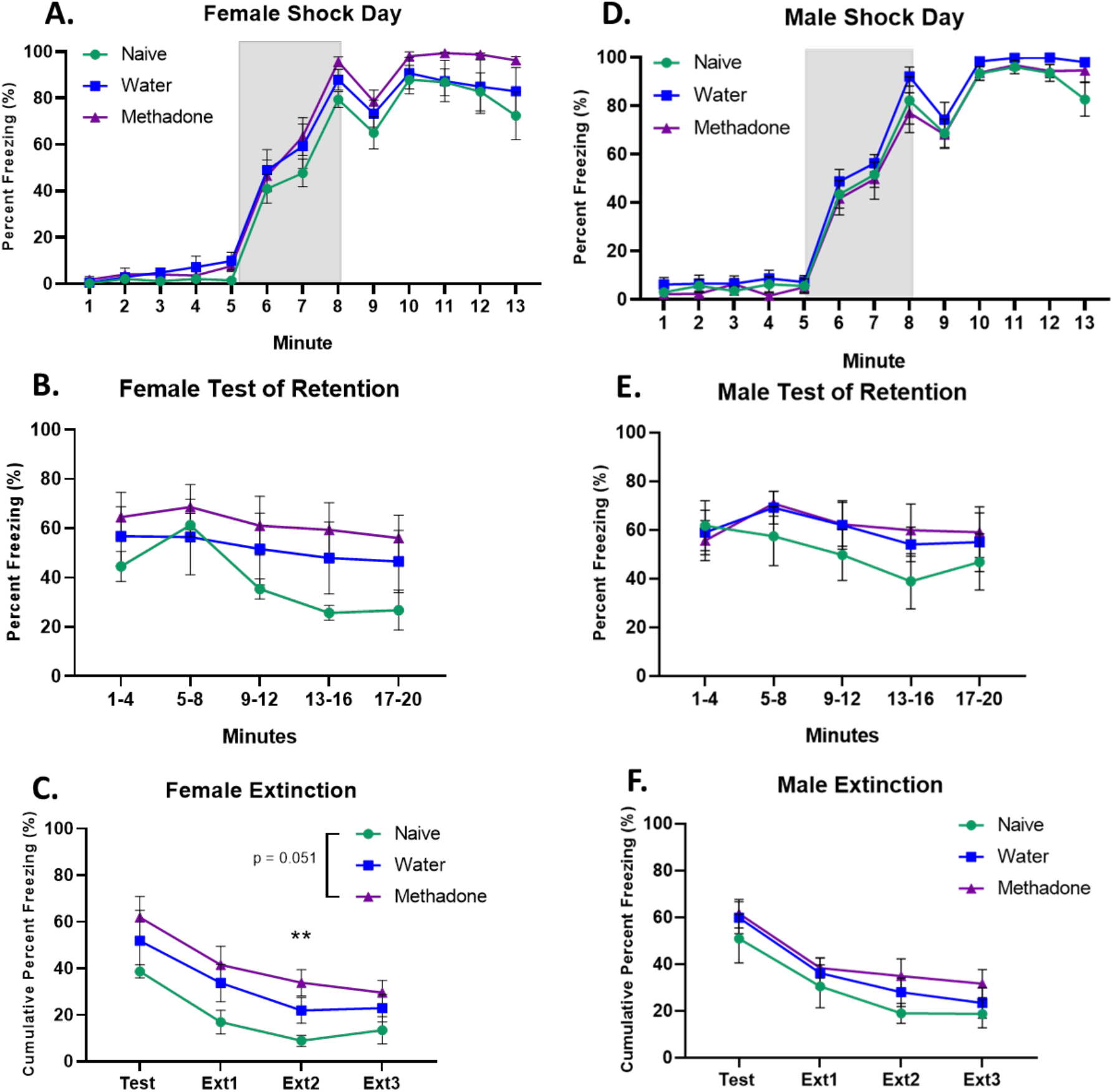
PME heightens female freezing during extinction learning. PME did not significantly alter percent freezing during the shock day or test of retention in females (A-B) or males (D-E). However, PME females showed significantly more freezing during extinction compared to naïve controls, particularly on extinction day 2 (C), but PME males did not show any differences in extinction (F). Grey box (A, D) represents timeframe when 3 shocks were delivered. Retention data (B, E) were analyzed across full 20min session but binned into 4 min periods in the figure for ease of visualization. Data points represent values from individual animals; error bars represent SEM. ** p < 0.01 compared to naïve.

For extinction learning, females showed a significant difference in exposure across testing days (exposure: *F*(2,21) = 3.447, *p* = 0.051; day: *F*(1.977, 41.53) = 35.37, *p* < 0.0001, no significant interaction: *F*(6,63) = 0.325, *p* = 0.921), with post hoc analysis showing this was driven by significantly more freezing in the methadone-exposed females compared to naïve females on the second extinction day (*p* = 0.006), but no significant difference in the water-exposed offspring compared to navie (*p* = 0.128) or methadone (*p* = 0.773) **(Fig. 5C).**

Males also exhibited typical conditioning behavior on the shock day, not showing any increases in percent freezing during the initial acclimation period, followed by substantial increases once the foot shocks began and persisting into the 5min consolidation period (time: (*F* (4.357, 91.49) = 335.600, *p* < 0.0001). However, there was no effect of exposure on percent freezing (*F*(2, 21) = 1.809, *p* = 0.189) and no significant interaction (*F*(24, 252) = 0.518, *p* = 0.971) **(Fig. 5D)**. On the test day, males still showed a significant main effect of time, decreasing freezing across the 20min test period (*F*(5.639, 118.4) = 3.715, *p* = 0.003), but no significant differences across exposure groups (*F*(2, 21) = 0.503, *p* = 0.612) and no interaction (*F*(38, 399) = 1.061, *p* = 0.376) **(Fig. 5E).** Finally, males did show significant decreases in freezing across extinction days (*F*(2.637, 55.38) = 43.2, *p* < 0.0001) but no significant differences across exposure groups for extinction days (*F*(2,21) = 0.999, *p* = 0.385) and no significant interaction (*F*(6,63) = 0.303*, p* = 0.9333) (**Fig. 5F).**

### 3.4 PME does not impair proliferation in the adult hippocampus of either sex

As discussed, one group of BrdU-injected offspring were sacrificed 24 hr after the final injection for assessment of dentate granule cell proliferation. Slices containing the dentate gyrus of the dHPC were double stained for BrdU and GFAP using immunofluorescence (**Fig. 9A)**. Cell counts represent the total number of cells with each phenotype summed from both hemispheres of six slices per animal. In females, there were no significant differences across exposure groups for BrdU^+^ cell counts (*F*(2,23) = 2.503, *p* = 0.104), GFAP^+^ cell counts (*F*(2,23) = 0.937, *p* = 0.406), or colabeled (BrdU^+^/GFAP^+^) (*F*(2,23) = 2.682, *p* = 0.09) **(Fig. 6A-C).** Male offspring showed similar results, with no significant differences across exposure in BrdU^+^ cell counts (*F*(2,21) = 0.152, *p* = 0.860), GFAP^+^ cell counts (*F*(2,21) = 0.081, *p* = 0.922), or colabeled (BrdU^+^/GFAP^+^) (*F*(2,21) = 0.098, *p* = 0.907) **(Fig. 6D-F).**

**Figure 6.**
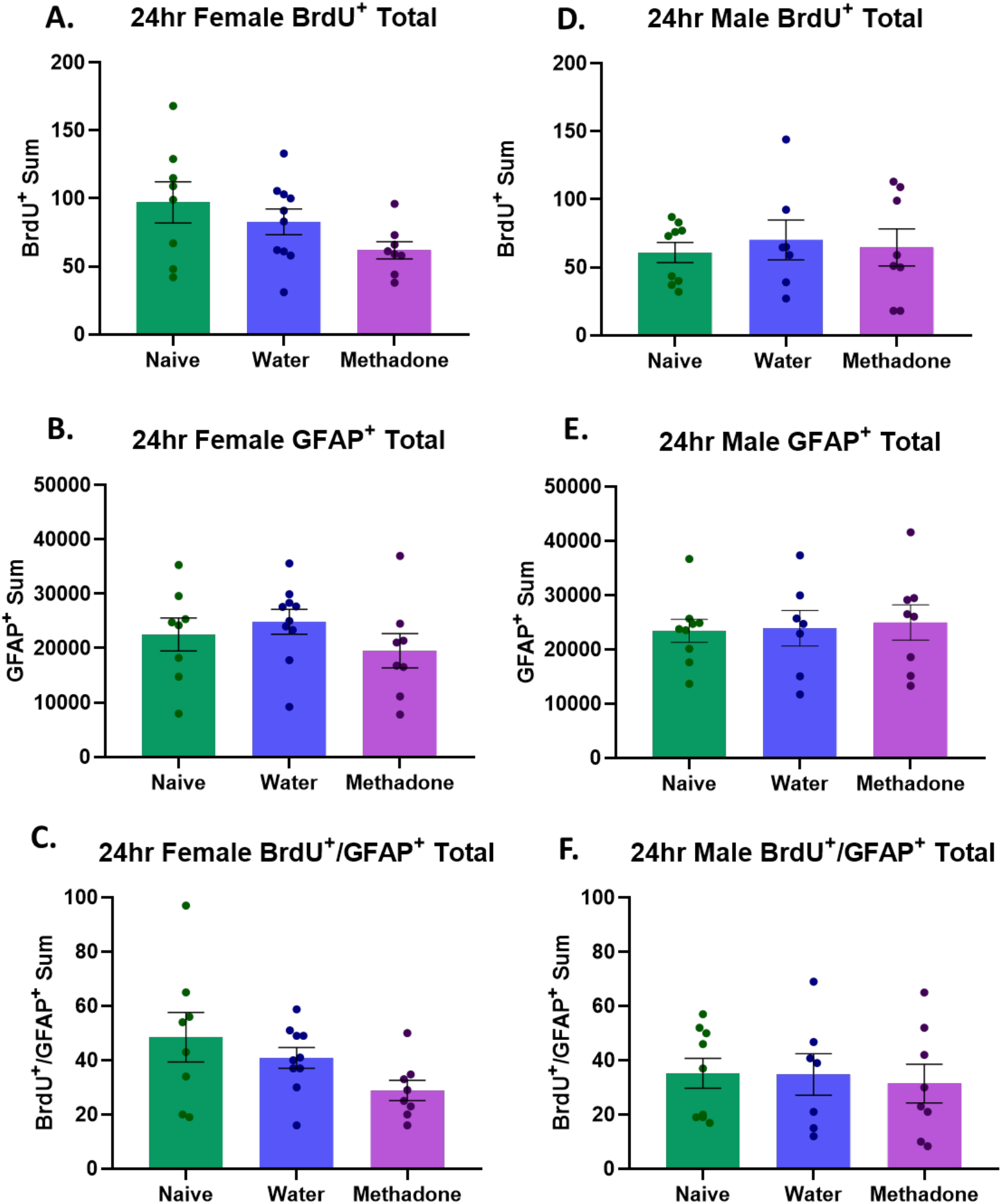
PME does not impair adult hippocampal proliferation in either sex. At the 24hr timepoint, female offspring showed no significant differences in the total number of BrdU^+^ cells (A), GFAP^+^ cells (B), or colabeled cells (C). Males also showed no significant differences in total numbers of BrdU^+^ cells (D), GFAP^+^ cells (E), or colabeled cells (F). Sums represent total counts of each phenotype for both hemispheres of 6 slices from the dorsal hippocampus. Data points represent values from individual animals; error bars represent SEM. BrdU = bromodeoxyuridine, GFAP = glial fibrillary acidic protein

Colabeled data were also analyzed as percent of BrdU^+^ that were colabeled with GFAP, but neither sex showed any significant differences across exposure (female: *F*(2.23) = 0.517, *p* = 0.603; males: *F*(2,21) = 0.948, *p* = 0.403; **Supplementary Fig. 4).** Interestingly, at this timepoint, female naïve offspring showed significantly more BrdU^+^ cells than male naïve offspring (*t*(15) = 2.224, *p* = 0.042), indicating naïve females have increased proliferation compared to naïve males. However, there were no significant differences in naïve male and female GFAP^+^ cell counts (*t*(15) = 0.246, *p* = 0.809) **(Supplementary Fig. 4).**

### 3.5 PME impairs female adult-born neuron survival at an immature timepoint

To examine differentiation and survival of adult-born neurons to an immature phenotype, a separate group of offspring were sacrificed 2 weeks after BrdU injections and stained, as described above, for BrdU and DCX (**Fig. 9B**). In females, PME offspring showed a significant reduction in BrdU^+^ cell counts compared to water and naïve offspring (*F*(2,23) = 3.347, *p* = 0.053; **Fig. 7A**). However, there were no significant differences among exposure groups for the number of DCX^+^ cells (*F*(2,23) = 1.889, *p* = 0.174) or number of BrdU^+^/DCX^+^ cells (*H*(2) = 4.903, *p* = 0.086) **(Fig. 7B-C)**. For males, there were no significant differences across exposure groups for BrdU^+^ cell counts (*F*(2,21) = 2.555, *p* = 0.102), DCX^+^ cell counts (*F*(2,21) = 2.456, *p* = 0.110), or number of colabeled BrdU^+^/DCX^+^ cells (*F*(2,21) = 1.599, *p* = 0.226) **(Fig. 7D-F).** Similar to above, we also analyzed data for percent of BrdU^+^ that were also colabeled with DCX. Interestingly, in females, naïve offspring did show significantly more percent colabeled with DCX (*F*(2.23) = 4.248, *p* = 0.027), with post hoc tests revealing this driven by a reduction in the water-exposed offspring compared to naïve (*p* = 0.026), but no difference between naïve and methadone-exposed (*p* = 0.154) or water- and methadone-exposed (*p* = 0.776). Male offspring did not show any significant difference in the number of BrdU^+^ colabeled with DCX (*F*(2.21) = 0.653, *p* = 0.531) (**Supplementary Fig. 5)**. Finally, there were no observed sex differences in the number of BrdU^+^ cells in naïve male and female offspring (*t*(16) = 0.489, *p* = 0.631) and no significant differences in naïve male and female DCX^+^ counts (*t*(16) = 1.730, *p* = 0.103); Supplementary Fig. 5).

**Figure 7.**
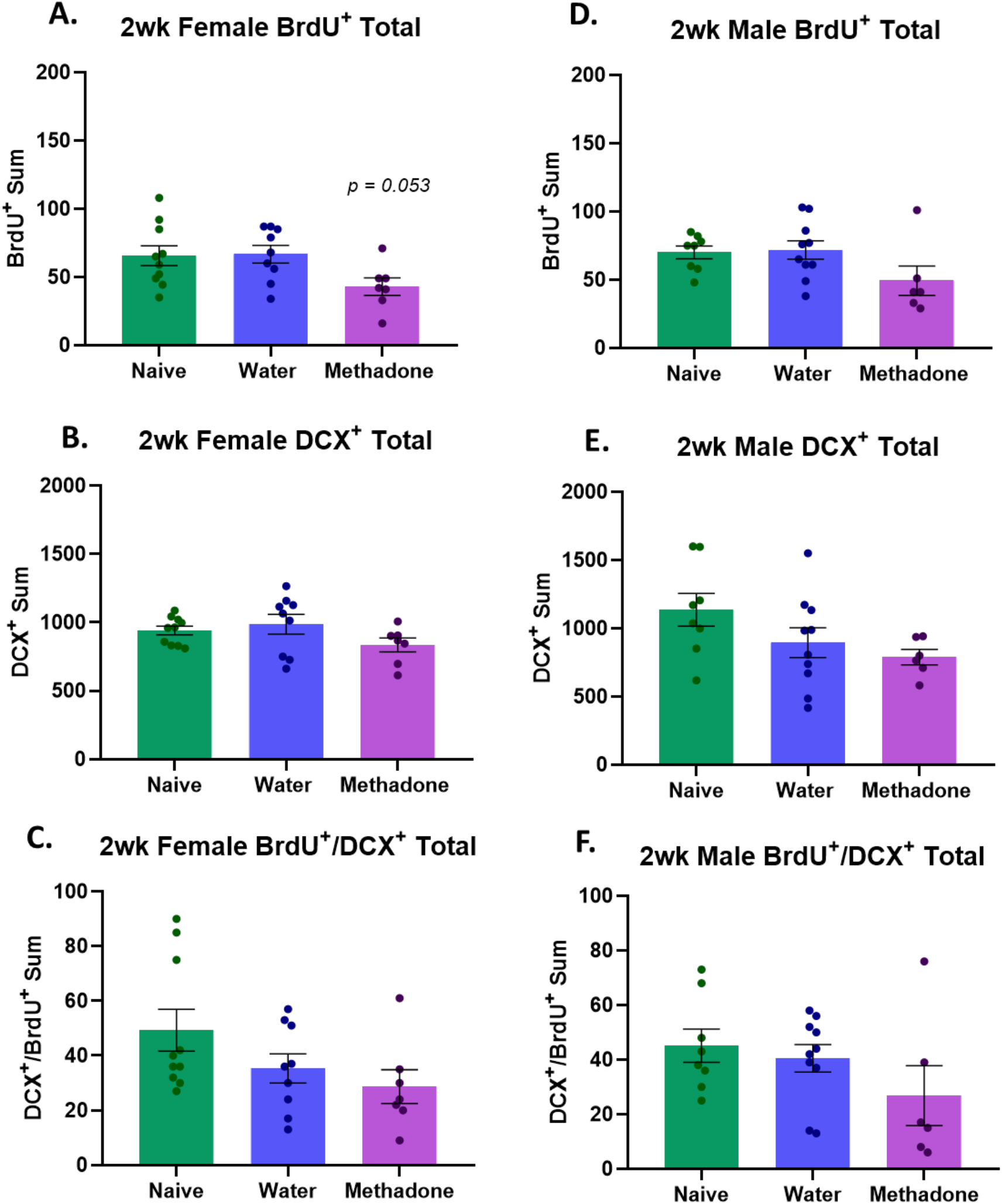
PME reduces the number of adult-born neurons at an immature timepoint in female offspring. PME females showed a reduced number of BrdU^+^ cells at the 2-week timepoint (A), but no differences in the number of DCX^+^ cells (B) or colabeled cells (C). Males did not show any significant differences in total numbers of BrdU^+^ cells (D), DCX^+^ cells (E), or colabeled cells (F). Sums represent total counts of each phenotype for both hemispheres of six slices from the dorsal hippocampus. Data points represent values from individual animals; error bars represent SEM. BrdU = bromodeoxyuridine, DCX = doublecortin

### 3.6 PME impairs adult-born neuron maturation and survival to a mature phenotype

Finally, a separate group of offspring were sacrificed 4 weeks after BrdU injections to assess the survival of the labeled adult-born neurons to a mature phenotype (**Fig. 9C**). In females, PME offspring showed a significant reduction in BrdU^+^ cells (*F*(2,21) = 8.479, *p* = 0.002), with post hoc analyses showing PME females had reduced BrdU cells compared to both the water controls (*p* = 0.005) and naïve controls (*p* = 0.005), but no difference between water and naïve groups (*p* = 0.997) **(Fig. 8A)**. Females did not show any significant differences in the number of NeuN^+^ cells overall (*F*(2,21) = 1.196, *p* = 0.322; **Fig. 8B)**, but did show a significant reduction in the number of BrdU^+^/NeuN^+^ cells (*H*(2) = 10.79, *p* = 0.005), with post hoc analyses showing significant reductions in PME offspring compared to both the water controls (*p* = 0.0099) and naïve controls (*p* = 0.018), but no difference between water and naïve groups (*p* > 0.999) **(Fig 8C).**

**Figure 8.**
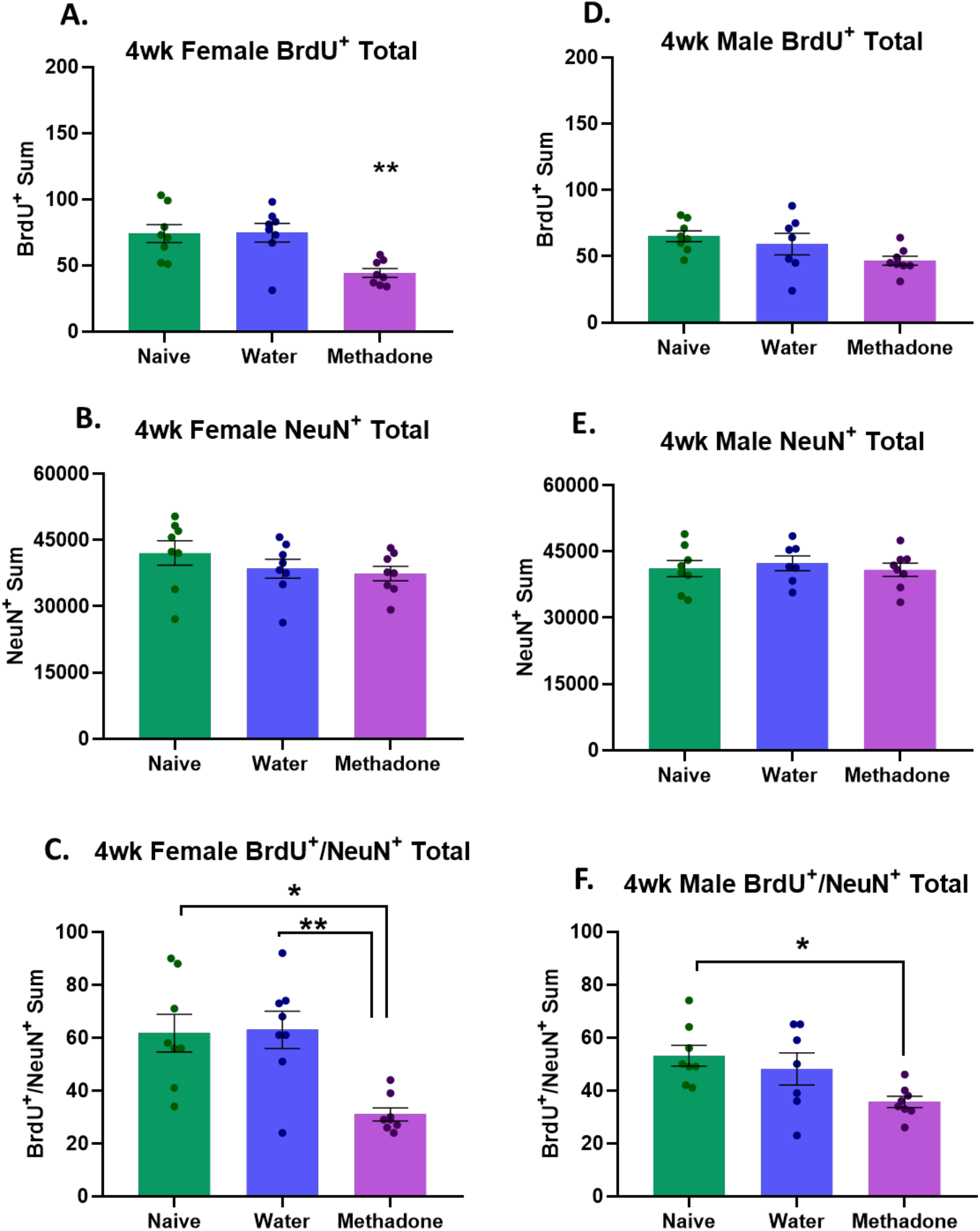
PME impairs adult-born neuronal survival. PME females showed a reduced number of BrdU^+^ cells (A) and colabeled cells (C) at the 4-week timepoint. However, there were no significant differences in the number of NeuN^+^ cells (B). Males showed a trend (*p* = 0.061) toward reduced total number of BrdU^+^ cells (D) and significant reductions in the number of colabeled cells (F), with no difference in the number of NeuN^+^ cells (E). Sums represent total counts of each phenotype for both hemispheres of 6 slices from the dorsal hippocampus. Data points represent values from individual animals; error bars represent SEM. * p < 0.05, ** p < 0.01. BrdU = bromodeoxyuridine, NeuN = neuronal nuclei

**Figure 9.**
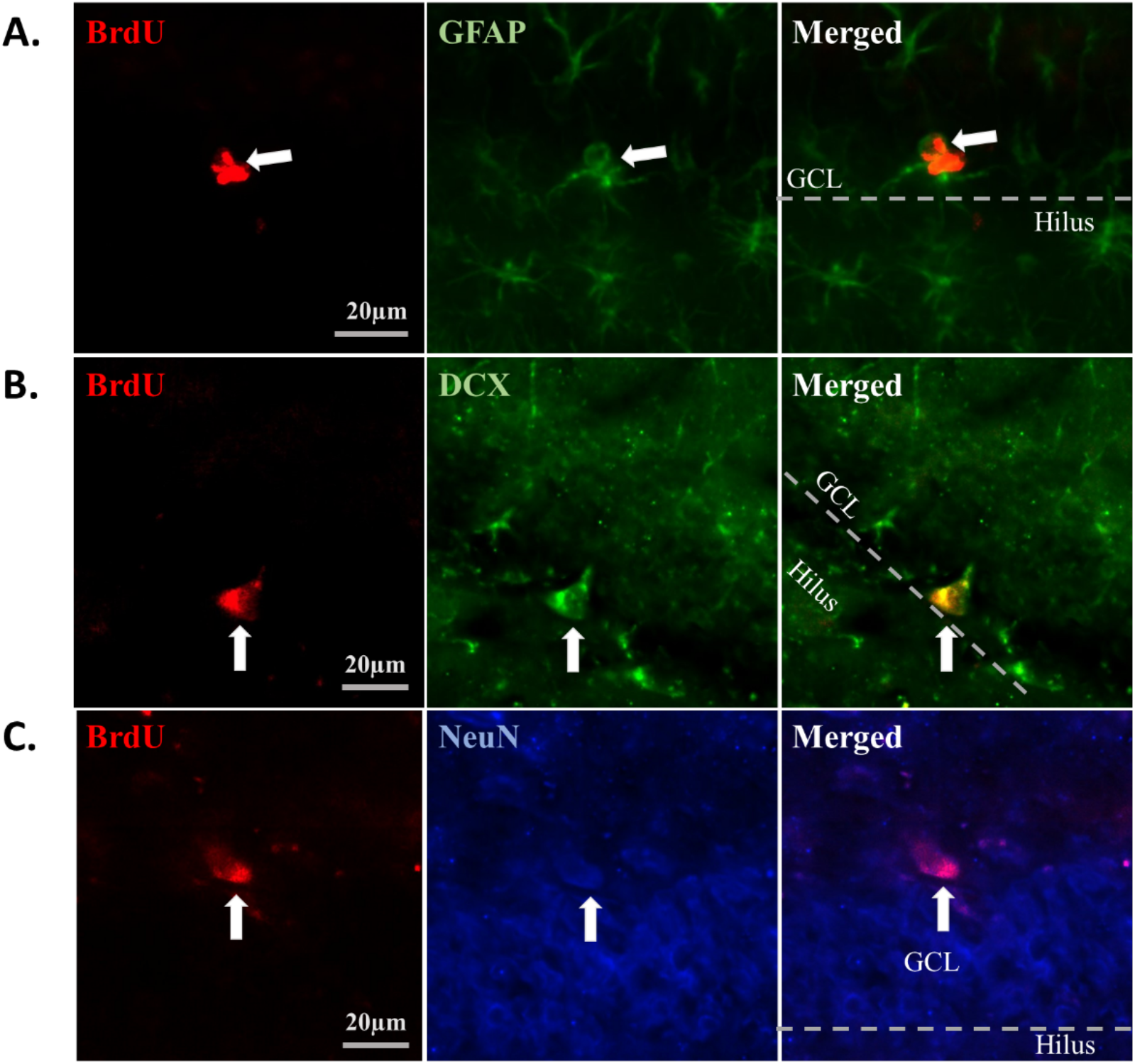
Representative images of immunofluorescent staining at 20X magnification. Each timepoint with their representative markers shown in each row. A.) 24hr staining for BrdU (red) and glial fibrillary acidic protein (GFAP; green), B.) 2-week staining for BrdU (red) and doublecortin (DCX; green), and C.) 4-week staining for BrdU (red) and neuronal nuclei (NeuN; blue). Colabeled cells are tracked through each row with white arrow. Dashed line shows approximate distinction between granule cell layer (GCL) and hilus of the dorsal hippocampus.

For male offspring, PME males showed a trend toward significant reductions in the number of BrdU^+^ cells (*F*(2,20) = 3.237, *p* = 0.061) **(Fig. 8D),** but no significant differences in the number of NeuN^+^ cells (*F*(2,20) = 0.208, *p* = 0.814) **(Fig. 8E)**. Importantly, males also showed a significant difference in the number of BrdU^+^/NeuN^+^ cells (*F*(2,20) = 4.809, *p* = 0.020), but post hoc tests showed that this was driven only by a reduction in the methadone-exposed offspring compared to naïve offspring (*p* = 0.018), with no difference between methadone and water groups (*p* = 0.119) or water and naïve control groups (*p* = 0.688) **(Fig. 8F).**

In analyses of the percent of BrdU^+^ cells also colabeled with NeuN, only female offspring showed a significant reduction in methadone-exposed animals (*F*(2,21) = 9.334, *p* = 0.001), with post hoc tests showing this was driven by reduction in methadone-exposed compared to both the water-exposed (*p* = 0.004) and naïve controls (*p* = 0.003), with no difference between water and naïve groups (*p* = 0.998). However, there were no differences in male offspring (*F*(2,20) = 0.890, *p*= 0.426) **(Supplementary Fig. 6)**. There were also no significant sex differences in naïve offspring for the number of BrdU^+^ cells (*t*(14) = 1.114, *p* = 0.284) or the number of NeuN^+^ cells (*t*(14) = 0.289, *p* = 0.777) **(Supplementary Fig. 6).**

## 4. Discussion

The present study aimed to continue characterizing the long-term learning and memory behaviors of offspring following PME, as well as assess the proliferation and survival of adult-born neurons in the dentate gyrus of the hippocampus in both sexes. Behaviorally, we have shown that adult PME offspring do not show impairments in spatial learning or retention using the Morris water maze, minimal disruptions in female pattern separation using the NOP task, and heightened freezing in adult females during the extinction period in a contextual fear conditioning paradigm. Additionally, we have shown that while both male and female offspring produce new DGCs at the same rate as control offspring, there are impairments in the number of these cells that are surviving to a mature neuron phenotype, 4 weeks later, with female offspring showing significant impairments at this timepoint and male offspring showing a trend toward the same. This is emphasized by a reduced number of the adult-born neurons that survive to this timepoint showing colabeling with the mature neuronal marker NeuN, indicating a possible delay in maturation of these cells in both sexes, which will be further discussed below.

Combined with the data presented in our previous work [28], this research adds a more complete picture to the current knowledge of the long-term effects of methadone exposure during pregnancy on the offspring. Overall, we have presented impairments in both behavior and hippocampal function, suggesting that female offspring may be particularly sensitive to these consequences.

As discussed previously, there are contradictory reports on the long-term effects of different POEs on spatial learning and memory in the Morris water maze. Many studies using different opioid exposures, like prenatal morphine or oxycodone exposure, have found delayed learning in the early acquisition phases of adult offspring [5, 42], in support of previous studies that showed similar effects in prenatal morphine -exposed adolescent offspring [22, 43].

However, those that examine the impacts of prenatal MOUDs on this behavior have not always shown these effects. Kongstorp and colleagues showed prenatal methadone, but not buprenorphine, produces impaired spatial memory in this task, but only on the 7-day retention test, with no differences in acquisition or short-term retention in P70 offspring [26]. This study also only conducted four training days for acquisition testing [26]. Additional studies have shown no changes in spatial learning or memory following these exposures, one with no age provided and only in male offspring [25] and one in weanling-age offspring prenatally exposed to buprenorphine [24]. Taken together, our findings support the literature that shows prenatal exposure to MOUDs may not produce the spatial impairments to the same extent as commonly misused opioids, as modeled with the oxycodone and morphine exposure studies. However, it is also important to note that related studies have varying results depending on specific procedural decisions, including the intertrial interval [42] and temperature of the water [35], so differential effects could also be explained by procedural differences and should be interpreted with this in mind.

Importantly, however, the contradictions in the Morris water maze literature expand beyond the context of POE, and into the adult neurogenesis field, as well. While there is substantial evidence showing that spatial learning and memory in rodents is dependent on adult-born neurons in the dentate gyrus [44–48], the intricacies of procedural differences mentioned above are also evident in the role of adult-born neurons in this task [for review, see [47]]. For example, some evidence suggests that adult hippocampal neurogenesis is heavily involved in reversal learning and cognitive flexibility assessments in this paradigm [47, 49] and the type of search strategy they use [46, 50], while classic parameters like latency and distance traveled may be less representative of changes in adult hippocampal neurogenesis [46, 47, 50]. Taken together, all of these factors may explain why we did not see any significant differences using our Morris water maze procedures, while there were still observed changes in adult hippocampal neurogenesis.

Regarding the NOP task, we have previously discussed the lack of literature relating to this behavior in the context of POE. This was an important behavior to assess, as rodent adult hippocampal neurogenesis is heavily involved in pattern separation abilities [38, 51–55]. Here, we have shown that female offspring prenatally exposed to both water and methadone do not differ from chance on the discrimination ratio, while naïve offspring investigated the object in the novel location significantly more than chance, as expected. The water-exposed females showed a slight trend toward increased investigation of the object in the novel location compared to chance (*p* = 0.078), where the methadone-exposed did not. However, overall, we cannot conclude that the lack of investigation of the object in the novel location is a methadone-specific impairment and may be the result of the procedural stress on the dams during pregnancy that both groups underwent. However, when analyzed as percent change from naïve offspring, PME females do show a significant reduction, while water-exposed do not show any significant differences compared to naïve offspring. All of this points to some subtle impairments in the PME females regarding pattern separation. Male offspring showed a few trends toward significant reduction in the PME offspring compared to control groups, but nothing significant.

This is consistent with a previous study that showed male pattern separation was intact following prenatal heroin exposure [27]. Together these findings suggest that PME does not have major effects on pattern separation in adult offspring; however, it is hard to make a conclusion about these effects without more POE studies to compare to. An important limitation that has been discussed previously [28] is that the naïve males did not differ from chance in the NOP task, as we would expect for typical pattern discrimination abilities of the controls. In this case, it is less complicated to interpret, since none of the three groups significantly differed from chance; however, it is still a limitation of the behavioral paradigm that the naïve controls did not behave as expected.

Finally, we did observe a significant increase in PME female freezing during the extinction learning period, with no differences in male offspring. This contrasts the one related study showing an impairment in male acquisition and extinction learning following prenatal morphine exposure [8]. However, this may point to males being less sensitive to PME compared to prenatal morphine exposure. Females were not included in the study by Tan and colleagues, so our findings are novel in showing alterations in fear responding in adulthood following PME, with very little information on female behavior in this task following other POEs. Additionally, previous work has shown differential responding to fear conditioning tasks in males and females, with males more often showing freezing behavior and females responding with active darting behavior [56]. This may contribute to not seeing differences in female freezing on shock day or during the retention test, but only in the assessments of extinction learning. This may also point to impairments not in the learning or memory for this task, but potentially in inhibition of fear responding, which may be driven by increases in anxiety-like behavior. This is important because it points to consequences in regulating fear responses or reducing fear in response to a stimulus that may not be threatening. Translationally, this could be viewed as the inability to inhibit a fear response, as is common in anxiety disorders and post-traumatic stress disorder, and may limit the effectiveness of behavioral therapies [57, 58]. While the initial phases of contextual fear conditioning (acquisition and retention test) are commonly linked with adult hippocampal neurogenesis [47, 59], neurogenesis has also been shown to play a vital role in fear memory extinction [47, 60–62] and fear generalization [63].

The adult hippocampal neurogenesis findings presented here are a novel addition to the understanding of hippocampal dysfunction in the adult brain following POEs, and specifically PME. In females we have shown no differences in proliferation when assessed at the 24hr timepoint, significant reductions in adult-born neurons at both the 2wk and 4wk timepoints, and a reduction in the number of adult-born cells labeled with the mature phenotype marker, NeuN, at the 4wk timepoint. This tells us that PME may not suppress proliferation of adult-generated cells in this region, but instead, lead to impairments relating to the survival and maturation of these neurons to a final mature phenotype. This would lead to a reduced number of mature granule cells that are integrating and properly functioning in the hippocampal circuit, which can explain some of the hippocampal-dependent behavioral deficits we have observed in PME females. It is important to note that these reductions in females were only present in the methadone-exposed offspring, not the water-exposed, which tells us that these impairments are not attributed to any stress-related effects of the prenatal procedures used, but specific to methadone.

This, however, was not the case for the male offspring, who showed a reduction in the number of adult-born neurons colabeled with NeuN at the 4wk timepoint, but only in the methadone-exposed compared to naïve controls. However, the males appear to be more sensitive to the prenatal stress effects of the procedures, as the water-exposed groups showed no significant difference to either group, falling in the middle of the two. Previous work has shown significant adult hippocampal impairments, including those relating to neurogenesis, as a result of prenatal stress exposure in both sexes [64] and when only looking at males [65], with some evidence showing male offspring may be more sensitive to the prenatal stress-induced effects on hippocampal neurogenesis and related behaviors [66]. However, early studies only looked in male offspring, so it is hard to determine if this really is sex-dependent. Nonetheless, the contribution of prenatal stress effects in these findings should be considered where naïve and water-exposed control group results differ. Additionally, PME males did not show a significant reduction in the number of adult-born neurons at this 4 wk timepoint, only a trend.

Unfortunately, in the PME male group at this timepoint, two brains were lost to unavoidable procedural events in the staining process, so this group is underpowered at *n* = 6 and may lack significance without the full power for the analyses.

Additional studies are required to fully assess this effect of delayed maturation and impaired survival of adult-born neurons following PME. For example, Yagi and colleagues have recently shown that there are sex differences in the maturation of these neurons, and that peak timepoints for the additional markers (DCX and NeuN) can vary across the time course of neuronal development [41]. The mature timepoint we selected, 4 weeks, may have been on the early side of the developmental window, as previous work has shown it takes approximately 4-8 weeks for DCGs to reach a mature phenotype [67, 68]. This means that for the neurons that were not colabeled with NeuN at the 4 wk timepoint, they may not yet have reached this mature phenotype, and brain collection a few weeks later would have shown higher percentages of mature BrdU^+^ neurons. However, this conclusion is not something we can fully support with the present data, and further research will need to follow up with a longer time frame and more frequent timepoints to identify any changes in the time course of maturation of these adult-born DGCs in PME offspring, as well as how it may differ by sex in the context of PME. Importantly, additional studies could examine these effects in combination with the previously mentioned studies that have shown reductions in hippocampal BDNF following POEs [12, 13, 69], particularly in females [5]. This is important because, as discussed, BDNF is a major regulator of neurogenesis, and particularly the survival of these adult-born neurons [70, 71].

Reduced activity or dysfunction of this protein may help to explain the reduction in survival, but not proliferation, that we saw in the current study. However, there are numerous other factors and mechanisms that contribute to these cells’ health and survival, including GABA function in the hippocampus, which we have previously shown is disrupted in adult female offspring following PME [28], and future work should further assess what mechanisms may ultimately be leading to these reductions.

Finally, some additional limitations to note about the current study. In previous work [28], we have thoroughly discussed the limitation of our PME model, and those also apply here, as the same PME procedures were used. These includes limitations to translatability of the model and the additive effects of prenatal stress that were discussed above. In the current study, there were also several limitations regarding the neurogenesis experiments. First, our goal was to have *n* = 8-10 per group, per sex, per time point. However, due to the numerous steps in this experimental timeline, there were many opportunities for loss of sample and we ended up with *n* = 6-7 in some groups. These were all unavoidable, but due to the typical variability in this type of histological assessment, having 1 or 2 animals less in each group may have impacted the results. Additionally, performing 4 i.p. injections in one day for each animal was likely a stressor of its own, and stress can substantially impact the processes of adult hippocampal neurogenesis [72, 73]. However, all animals from all groups went through these procedures, so it is not likely that this effect would explain any of the group differences seen unless there are differences in the sensitivity to stress on adult neurogenesis as a result of PME. This question was outside the scope of our studies and has not been investigated thus far but should be followed up on in future research. Finally, one other step that would have strengthened this study would have been to make a direct link between the behaviors and adult hippocampal neurogenesis. Our experiments were all in separate groups, and therefore, we cannot make correlations between the behavior and neurogenesis measures. Future work should continue to assess these cognitive and hippocampal disruptions following PME using measures that can make this direct correlation.

Taken together, this work has identified significant, long-term cognitive disruptions as a result of PME, and has continued to examine hippocampal mechanisms, namely dentate granule cells, that may be a target of this insult. Additionally, we continue to show that female offspring appear to be more sensitive to these effects as adults, which is an important consideration when continuing research in this field. This research is essential in helping to fully understand the impacts of opioid exposure during pregnancy, particularly MOUDs, as they are increasing in access and use for pregnant individuals. This knowledge will allow us to prepare and provide support for exposed individuals as they begin school and progress through the rest of their lives.

## Acknowledgements

We would like to thank Dr. Lisa Savage, Dr. Anushree Karkhanis, and Dr. Anna Klintsova, for their guidance in the development of this project, as well as Dr. Molly Deak and Nicole Reitz for their time in training of required techniques. We would also like to thank Dana Silberstein for assisting in exposures for this work. This work was funded by the Binghamton University Department of Psychology and Harpur Faculty Research Grant.

## References

1. Simmons, S.C., et al., Effects of prenatal opioid exposure on synaptic adaptations and behaviors across development. Neuropharmacology, 2022: p. 109312.

2. Drake, C.T., C. Chavkin, and T.A. Milner, Opioid systems in the dentate gyrus. Prog Brain Res, 2007. 163: p. 245–63.

3. Simmons, M.L. and C. Chavkin, Endogenous opioid regulation of hippocampal function. Int Rev Neurobiol, 1996. 39: p. 145–96.

4. Stumm, R.K., et al., Neuronal types expressing mu- and delta-opioid receptor mRNA in the rat hippocampal formation. J Comp Neurol, 2004. 469(1): p. 107–18.

5. Ahmadalipour, A., et al., Deleterious effects of prenatal exposure to morphine on the spatial learning and hippocampal BDNF and long-term potentiation in juvenile rats: Beneficial influences of postnatal treadmill exercise and enriched environment. Neurobiol Learn Mem, 2018. 147: p. 54–64.

6. Lin, C.S., et al., Prenatal morphine alters the synaptic complex of postsynaptic density 95 with N-methyl-D-aspartate receptor subunit in hippocampal CA1 subregion of rat offspring leading to long-term cognitive deficits. Neuroscience, 2009. 158(4): p. 1326–37.

7. Niu, L., et al., Impaired in vivo synaptic plasticity in dentate gyrus and spatial memory in juvenile rats induced by prenatal morphine exposure. Hippocampus, 2009. 19(7): p. 649–57.

8. Tan, J.W., et al., Impaired contextual fear extinction and hippocampal synaptic plasticity in adult rats induced by prenatal morphine exposure. Addict Biol, 2015. 20(4): p. 652–62.

9. Velisek, L., R. Slamberova, and I. Vathy, Prenatal morphine exposure suppresses mineralocorticoid receptor-dependent basal synaptic transmission and synaptic plasticity in the lateral perforant path in adult male rats. Brain Res Bull, 2003. 61(6): p. 571–6.

10. Villarreal, D.M., B. Derrick, and I. Vathy, Prenatal morphine exposure attenuates the maintenance of late LTP in lateral perforant path projections to the dentate gyrus and the CA3 region in vivo. J Neurophysiol, 2008. 99(3): p. 1235–42.

11. Wu, C.C., et al., Prenatal buprenorphine exposure decreases neurogenesis in rats. Toxicol Lett, 2014. 225(1): p. 92–101.

12. Nasiraei-Moghadam, S., et al., Reversal of prenatal morphine exposure-induced memory deficit in male but not female rats. J Mol Neurosci, 2013. 50(1): p. 58–69.

13. Schrott, L.M., L. Franklin, and P.A. Serrano, Prenatal opiate exposure impairs radial arm maze performance and reduces levels of BDNF precursor following training. Brain Res, 2008. 1198: p. 132–40.

14. Bekinschtein, P., et al., Brain-derived neurotrophic factor interacts with adult-born immature cells in the dentate gyrus during consolidation of overlapping memories. Hippocampus, 2014. 24(8): p. 905–11.

15. Cirulli, F., et al., Intrahippocampal administration of BDNF in adult rats affects short-term behavioral plasticity in the Morris water maze and performance in the elevated plus-maze. Hippocampus, 2004. 14(7): p. 802–7.

16. Kesslak, J.P., et al., Learning upregulates brain-derived neurotrophic factor messenger ribonucleic acid: a mechanism to facilitate encoding and circuit maintenance? Behav Neurosci, 1998. 112(4): p. 1012–9.

17. Mizuno, M., et al., Involvement of brain-derived neurotrophic factor in spatial memory formation and maintenance in a radial arm maze test in rats. J Neurosci, 2000. 20(18): p. 7116–21.

18. Lum, J.S., et al., Prenatal methadone exposure impairs adolescent cognition and GABAergic neurodevelopment in a novel rat model of maternal methadone treatment. Prog Neuropsychopharmacol Biol Psychiatry, 2021. 110: p. 110281.

19. SAMHSA, Clinical Guidance for Treating Pregnant and Parenting Women With Opioid Use Disorder and Their Infants. 2018, United States Department of Health and Human Services: Rockville, MD.

20. SAMHSA, A collaborative approach to the treatment of pregnant women with opioid use disorders. 2016: Rockville, MD.

21. ACOG, Opioid Use and Opioid Use Disorder in Pregnancy. 2017, American College of Obstetricians and Gynecologists and American Society of Addiction Medicine.

22. Yang, S.N., et al., Alterations of postsynaptic density proteins in the hippocampus of rat offspring from the morphine-addicted mother: Beneficial effect of dextromethorphan. Hippocampus, 2006. 16(6): p. 521–30.

23. Sarkaki, A., et al., Sex-specific effects of developmental morphine exposure and rearing environments on hippocampal spatial memory. Int J Dev Neurosci, 2022.

24. Hung, C.J., et al., Depression-like effect of prenatal buprenorphine exposure in rats. PLoS One, 2013. 8(12): p. e82262.

25. Chiang, Y.C., et al., Beneficial effects of co-treatment with dextromethorphan on prenatally methadone-exposed offspring. J Biomed Sci, 2015. 22: p. 19.

26. Kongstorp, M., et al., Prenatal exposure to methadone or buprenorphine impairs cognitive performance in young adult rats. Drug Alcohol Depend, 2020. 212: p. 108008.

27. Lu, R., et al., Effects of prenatal cocaine and heroin exposure on neuronal dendrite morphogenesis and spatial recognition memory in mice. Neurosci Lett, 2012. 522(2): p. 128–33.

28. Gamble, M.E., R. Marfatia, and M.R. Diaz, Prenatal methadone exposure leads to long-term memory impairments and disruptions of dentate granule cell function in a sex-dependent manner. Addict Biol, 2022. 27(5): p. e13215.

29. Nunez, J., Morris Water Maze Experiment. J Vis Exp, 2008(19).

30. Vorhees, C.V. and M.T. Williams, Morris water maze: procedures for assessing spatial and related forms of learning and memory. Nat Protoc, 2006. 1(2): p. 848–58.

31. Paxinos, G. and C. Watson, Paxino’s and Watson’s The rat brain in stereotaxic coordinates. Seventh edition. ed. 2014, Amsterdam ; Boston: Elsevier/AP, Academic Press is an imprint of Elsevier. 1 volume (unpaged).

32. Colon, L.M. and A.M. Poulos, Contextual processing elicits sex differences in dorsal hippocampus activation following footshock and context fear retrieval. Behav Brain Res, 2020. 393: p. 112771.

33. Cost, K.T., et al., Sex differences in object-in-place memory of adult rats. Behav Neurosci, 2012. 126(3): p. 457–64.

34. Jonasson, Z., Meta-analysis of sex differences in rodent models of learning and memory: a review of behavioral and biological data. Neuroscience and Biobehavioral Reviews, 2005. 28(8): p. 811–825.

35. O’Leary, T.P., et al., Sex Differences in the Spatial Behavior Functions of Adult-Born Neurons in Rats. eNeuro, 2022. 9(3).

36. Saucier, D., et al., Female advantage for object location memory in peripersonal but not extrapersonal space. J Int Neuropsychol Soc, 2007. 13(4): p. 683–6.

37. Saucier, D.M., et al., Sex differences in object location memory and spatial navigation in Long-Evans rats. Anim Cogn, 2008. 11(1): p. 129–37.

38. Yagi, S., et al., Sex and strategy use matters for pattern separation, adult neurogenesis, and immediate early gene expression in the hippocampus. Hippocampus, 2016. 26(1): p. 87–101.

39. Barker, J.M. and L.A. Galea, Repeated estradiol administration alters different aspects of neurogenesis and cell death in the hippocampus of female, but not male, rats. Neuroscience, 2008. 152(4): p. 888–902.

40. Falconer, E.M. and L.A. Galea, Sex differences in cell proliferation, cell death and defensive behavior following acute predator odor stress in adult rats. Brain Res, 2003. 975(1-2): p. 22–36.

41. Yagi, S., et al., Sex Differences in Maturation and Attrition of Adult Neurogenesis in the Hippocampus. eNeuro, 2020. 7(4).

42. Davis, C.P., et al., Prenatal oxycodone exposure impairs spatial learning and/or memory in rats. Behav Brain Res, 2010. 212(1): p. 27–34.

43. Yang, S.N., et al., Prenatal administration of morphine decreases CREBSerine-133 phosphorylation and synaptic plasticity range mediated by glutamatergic transmission in the hippocampal CA1 area of cognitive-deficient rat offspring. Hippocampus, 2003. 13(8): p. 915–921.

44. Deng, W., et al., Adult-born hippocampal dentate granule cells undergoing maturation modulate learning and memory in the brain. J Neurosci, 2009. 29(43): p. 13532–42.

45. Dupret, D., et al., Spatial relational memory requires hippocampal adult neurogenesis. PLoS One, 2008. 3(4): p. e1959.

46. Garthe, A., J. Behr, and G. Kempermann, Adult-generated hippocampal neurons allow the flexible use of spatially precise learning strategies. PLoS One, 2009. 4(5): p. e5464.

47. Hernandez-Mercado, K. and A. Zepeda, Morris Water Maze and Contextual Fear Conditioning Tasks to Evaluate Cognitive Functions Associated With Adult Hippocampal Neurogenesis. Front Neurosci, 2021. 15: p. 782947.

48. Rola, R., et al., Radiation-induced impairment of hippocampal neurogenesis is associated with cognitive deficits in young mice. Exp Neurol, 2004. 188(2): p. 316–30.

49. Webler, R.D., et al., Maturational phase of hippocampal neurogenesis and cognitive flexibility. Neurosci Lett, 2019. 711: p. 134414.

50. Gil-Mohapel, J., et al., Hippocampal neurogenesis levels predict WATERMAZE search strategies in the aging brain. PLoS One, 2013. 8(9): p. e75125.

51. Abu, Y. and S. Roy, Prenatal opioid exposure and vulnerability to future substance use disorders in offspring. Exp Neurol, 2021. 339: p. 113621.

52. Clelland, C.D., et al., A functional role for adult hippocampal neurogenesis in spatial pattern separation. Science, 2009. 325(5937): p. 210–3.

53. Nakashiba, T., et al., Young dentate granule cells mediate pattern separation, whereas old granule cells facilitate pattern completion. Cell, 2012. 149(1): p. 188–201.

54. Sahay, A., et al., Increasing adult hippocampal neurogenesis is sufficient to improve pattern separation. Nature, 2011. 472(7344): p. 466–70.

55. Franca, T.F.A., et al., Hippocampal neurogenesis and pattern separation: A meta-analysis of behavioral data. Hippocampus, 2017. 27(9): p. 937–950.

56. Gruene, T.M., et al., Sexually divergent expression of active and passive conditioned fear responses in rats. Elife, 2015. 4.

57. Pittig, A., L. van den Berg, and B. Vervliet, The key role of extinction learning in anxiety disorders: behavioral strategies to enhance exposure-based treatments. Curr Opin Psychiatry, 2016. 29(1): p. 39–47.

58. Anderson, K.C. and T.R. Insel, The promise of extinction research for the prevention and treatment of anxiety disorders. Biol Psychiatry, 2006. 60(4): p. 319–21.

59. Besnard, A. and A. Sahay, Enhancing adult neurogenesis promotes contextual fear memory discrimination and activation of hippocampal-dorsolateral septal circuits. Behav Brain Res, 2021. 399: p. 112917.

60. Deng, W., J.B. Aimone, and F.H. Gage, New neurons and new memories: how does adult hippocampal neurogenesis affect learning and memory? Nat Rev Neurosci, 2010. 11(5): p. 339–50.

61. Pan, Y.W., et al., Inhibition of adult neurogenesis by inducible and targeted deletion of ERK5 mitogen-activated protein kinase specifically in adult neurogenic regions impairs contextual fear extinction and remote fear memory. J Neurosci, 2012. 32(19): p. 6444–55.

62. Pan, Y.W., D.R. Storm, and Z. Xia, Role of adult neurogenesis in hippocampus-dependent memory, contextual fear extinction and remote contextual memory: new insights from ERK5 MAP kinase. Neurobiol Learn Mem, 2013. 105: p. 81–92.

63. Besnard, A. and A. Sahay, Adult Hippocampal Neurogenesis, Fear Generalization, and Stress. Neuropsychopharmacology, 2016. 41(1): p. 24–44.

64. Mandyam, C.D., et al., Stress experienced in utero reduces sexual dichotomies in neurogenesis, microenvironment, and cell death in the adult rat hippocampus. Dev Neurobiol, 2008. 68(5): p. 575–89.

65. Lemaire, V., et al., Prenatal stress produces learning deficits associated with an inhibition of neurogenesis in the hippocampus. Proc Natl Acad Sci U S A, 2000. 97(20): p. 11032–7.

66. Weinstock, M., Gender differences in the effects of prenatal stress on brain development and behaviour. Neurochem Res, 2007. 32(10): p. 1730–40.

67. Goncalves, J.T., S.T. Schafer, and F.H. Gage, Adult Neurogenesis in the Hippocampus: From Stem Cells to Behavior. Cell, 2016. 167(4): p. 897–914.

68. Toda, T., et al., The role of adult hippocampal neurogenesis in brain health and disease. Mol Psychiatry, 2019. 24(1): p. 67–87.

69. Ahmadalipour, A., et al., Effects of environmental enrichment on behavioral deficits and alterations in hippocampal BDNF induced by prenatal exposure to morphine in juvenile rats. Neuroscience, 2015. 305: p. 372–83.

70. Choi, S.H., et al., Regulation of hippocampal progenitor cell survival, proliferation and dendritic development by BDNF. Mol Neurodegener, 2009. 4: p. 52.

71. Sairanen, M., et al., Brain-derived neurotrophic factor and antidepressant drugs have different but coordinated effects on neuronal turnover, proliferation, and survival in the adult dentate gyrus. J Neurosci, 2005. 25(5): p. 1089–94.

72. Schoenfeld, T.J. and E. Gould, Stress, stress hormones, and adult neurogenesis. Exp Neurol, 2012. 233(1): p. 12–21.

73. Warner-Schmidt, J.L. and R.S. Duman, Hippocampal neurogenesis: opposing effects of stress and antidepressant treatment. Hippocampus, 2006. 16(3): p. 239–49.

