## Supplementary Material for "Prenatal Methadone Exposure Leads To Disruptions In Adult-Born Dentate Granule Cell Survival And Female Persistent Fear Responding"

### Supplementary Materials

| Table 1. Litter Representation | | # animals (# litters), animals per litter | |
|---|---|---|---|
| **Experiment** | **Exposure** | **Male** | **Female** |
| Morris Water Maze | Naïve | n = 8(4), 2 per litter | n = 10(5), 2 per litter |
| | Water | n = 8(4), 2 per litter | n = 8(4), 2 per litter |
| | Methadone | n = 10(5), 2 per litter | n = 10(5), 2 per litter |
| Novel Object in Place | Naïve | n = 8(4), 2 per litter | n = 8(4), 2 per litter |
| | Water | n = 8(4), 2 per litter | n = 8(4), 2 per litter |
| | Methadone | n = 8(4), 2 per litter | n = 8(4), 2 per litter |
| Fear Conditioning | Naïve | n = 8(4), 2 per litter | n = 8(4), 2 per litter |
| | Water | n = 8(4), 2 per litter | n = 8(4), 2 per litter |
| | Methadone | n = 8(4), 2 per litter | n = 8(4), 2 per litter |
| Immunofluorescence (24hr) | Naïve | n = 9(6), 1-2 per litter | n = 8(5), 1-2 per litter |
| | Water | n = 7(5), 1-2 per litter | n = 10(5), 2 per litter |
| | Methadone | n = 8(6), 1-2 per litter | n = 8(5), 1-2 per litter |
| Immunofluorescence (2wk) | Naïve | n = 8(4), 2 per litter | n = 10(5), 2 per litter |
| | Water | n = 10(5), 2 per litter | n = 9(5), 1-2 per litter |
| | Methadone | n = 6(4), 1-2 per litter | n = 7(4), 1-2 per litter |
| Immunofluorescence (4wk) | Naïve | n = 8(4), 2 per litter | n = 8(4), 2 per litter |
| | Water | n = 7(4), 1-2 per litter | n = 8(4), 2 per litter |
| | Methadone | n = 8(4), 2 per litter | n = 8(4), 2 per litter |

**Supplementary Table 1.** Number of animals, litter representation, and animals per litter for each exposure group, sex, and experiment are displayed. n = number of animals (number of litters), number of animals per litter.

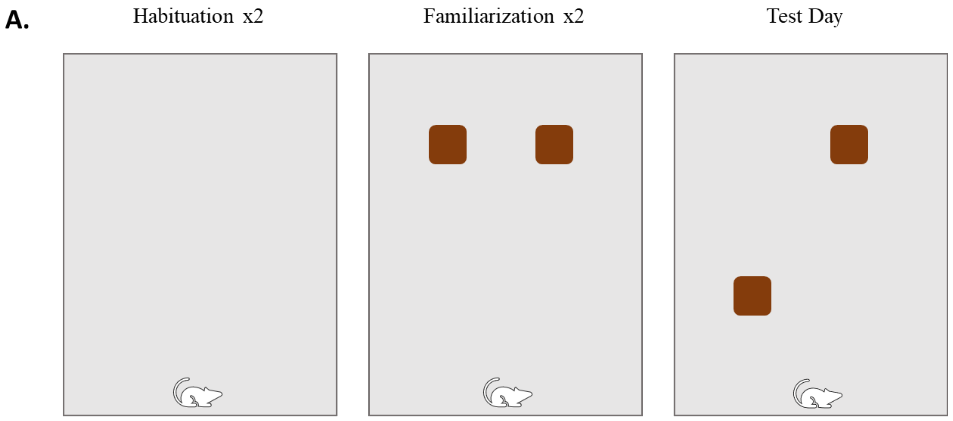


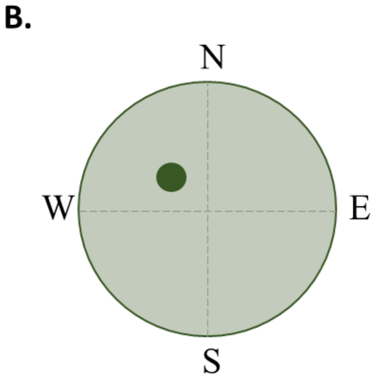


| Morris Water Maze Starting Positions | |
|---|---|
| **Day** | **Starting Positions (Trial 1, 2, 3, 4)** |
| Training Day 1 | S, E, NE, SW |
| Training Day 2 | NE, S, SW, N |
| Training Day 3 | SE, N, E, SW |
| Training Day 4 | S, NE, N, SE |
| Training Day 5 | N, SE, NE, SW |
| Training Day 6 | S, E, NE, SE |
| 24hr Test Day | SE |
| 7 day Test Day | SE |

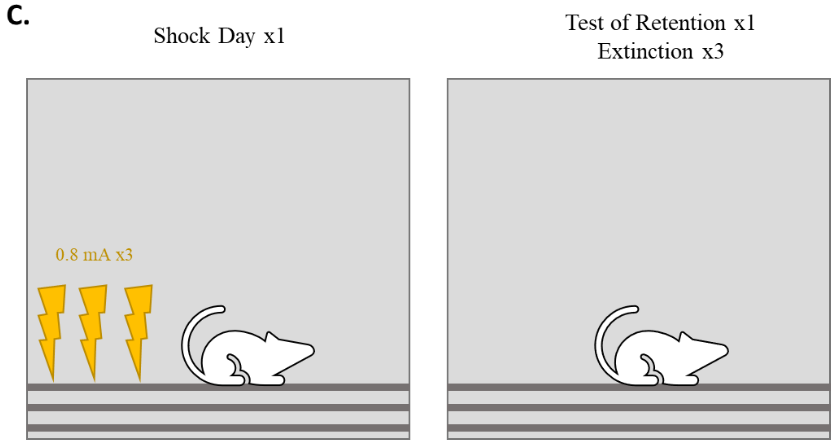


**Supplementary Figure 1.** Models demonstrating behavioral paradigm set up for the Novel Object in Place task (A), Morris Water maze pool and training positions (B), and Contextual Fear Conditioning (C).

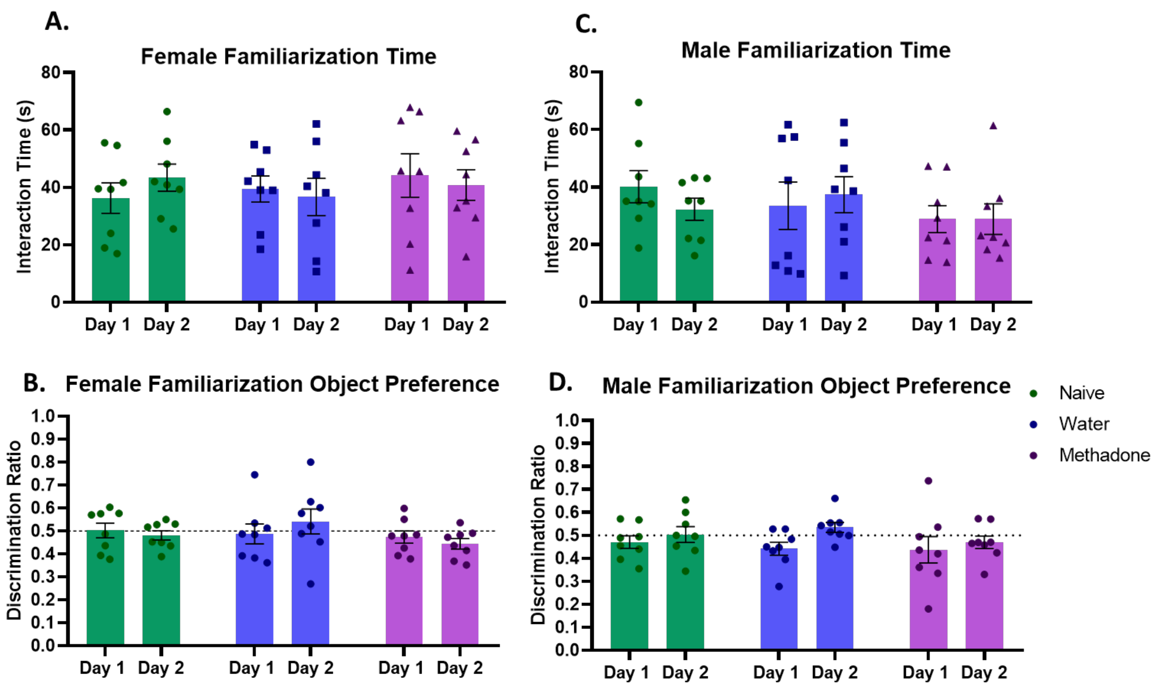


**Supplementary Figure 2.** Novel Object in Place training. There were no significant differences in total time spent investigating objects (A,C) and no side preferences (B,D) in either sex, across the three exposure groups for the 2 familiarization days of the Novel Object in Place task. Preference data are displayed as discrimination ratios for the left side object, with chance = 0.5. Interaction time displays the total time investigating either object during the session. These data indicate that any results found are not due to differences in encoding or preference for the object on one side. Data points represent individual animal scores; error bars represent SEM.

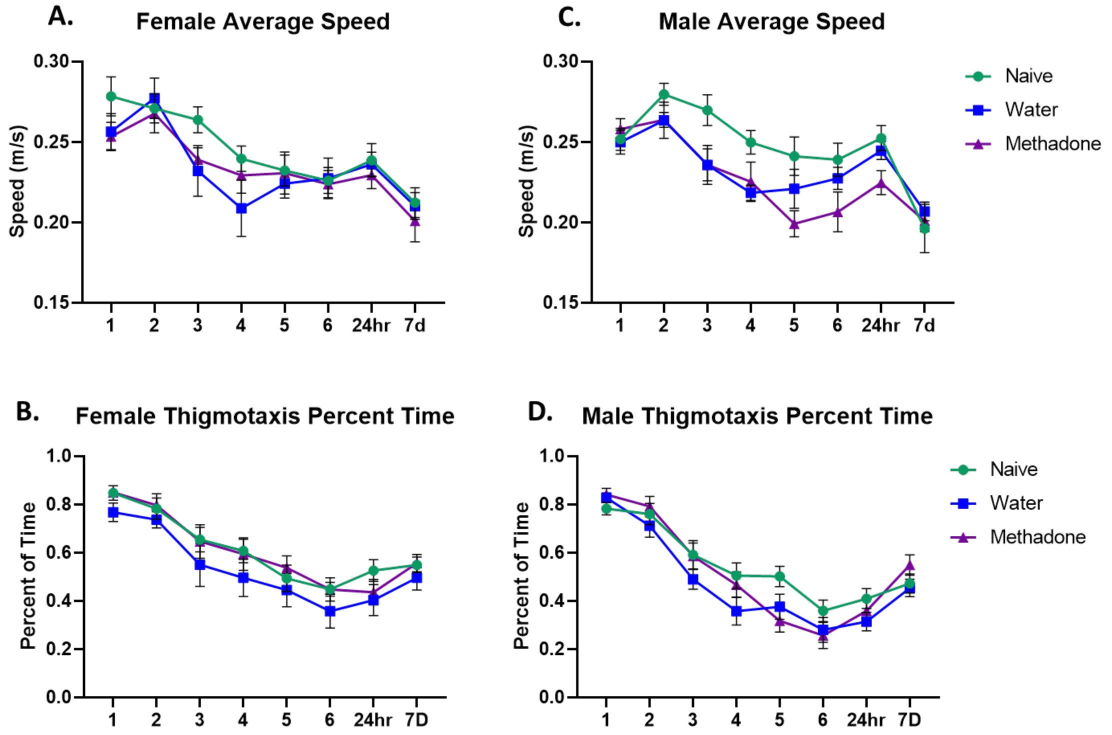


**Supplementary Figure 3.** Morris water maze extra measures. Females did not show any significant differences in average swimming speed (A) or thigmotaxis behavior (B). Males showed a significant interaction in average speed, driven by increased speed in naive males compared to water males on day 4 ($p = 0.009$) and increased speed in naïve controls compared to methadone on day 5 ($p = 0.031$) (C), but they did not show any significant differences in thigmotaxis behavior (D). Data point represent individual animal scores; error bars represent SEM.

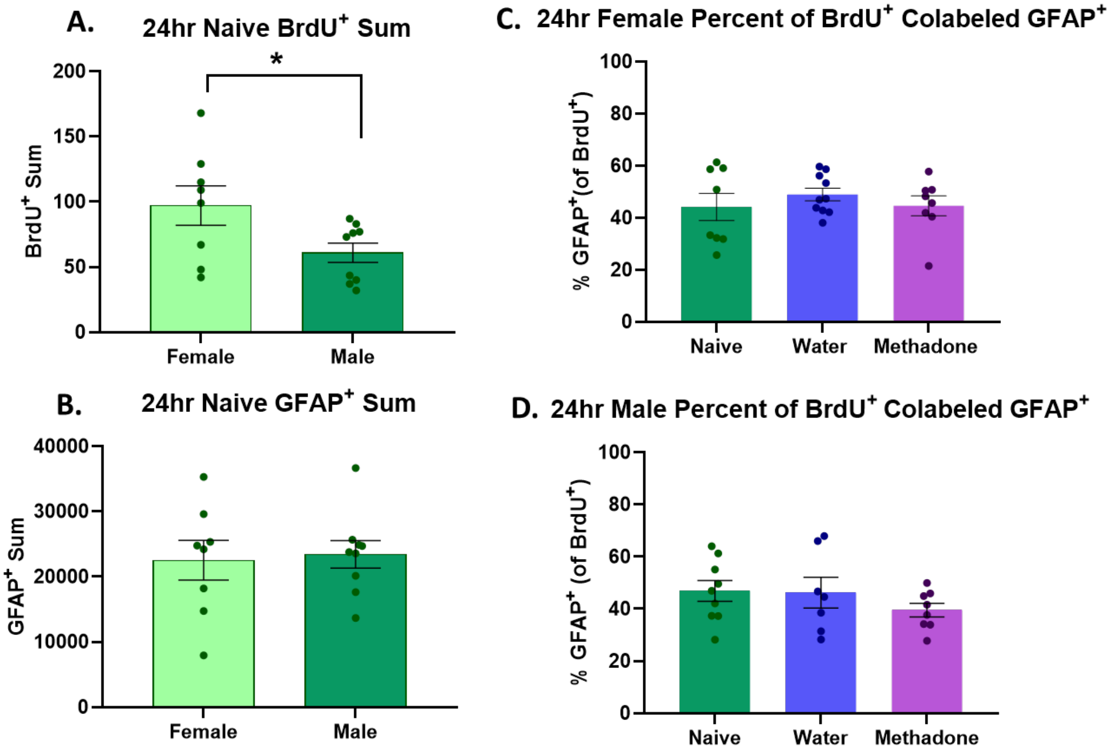


**Supplementary Figure 4.** Naïve females showed significantly more BrdU$^+$ cells than naïve males at the 24hr timepoint (A), but there were no sex differences in naïve animals for number of GFAP$^+$ cells (B). Neither sex showed a significant difference in the percent of BrdU$^+$ cells that were colabeled with GFAP (C-D). Data points represent individual animal counts; error bars represent SEM. * $p < 0.05$.

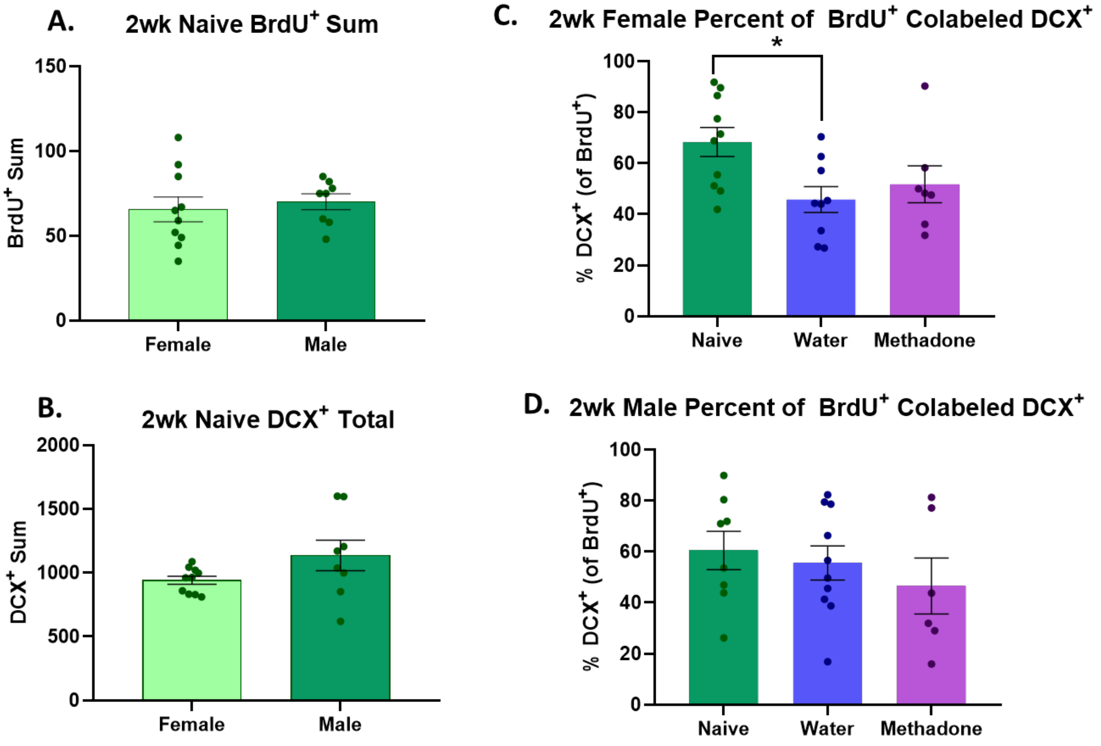


**Supplementary Figure 5.** There were no significant sex differences in the number of $BrdU^+$ cells in naïve controls at the 2wk timepoint (A), and no sex differences in naïve animals for number of $DCX^+$ cells (B). Water-exposed females showed a significant reduction in the number of $BrdU^+$ cells that were colabeled with DCX, but no difference in PME animals (C) and no difference in males (D). Data points represent individual animal counts; error bars represent SEM. * $p < 0.05$.

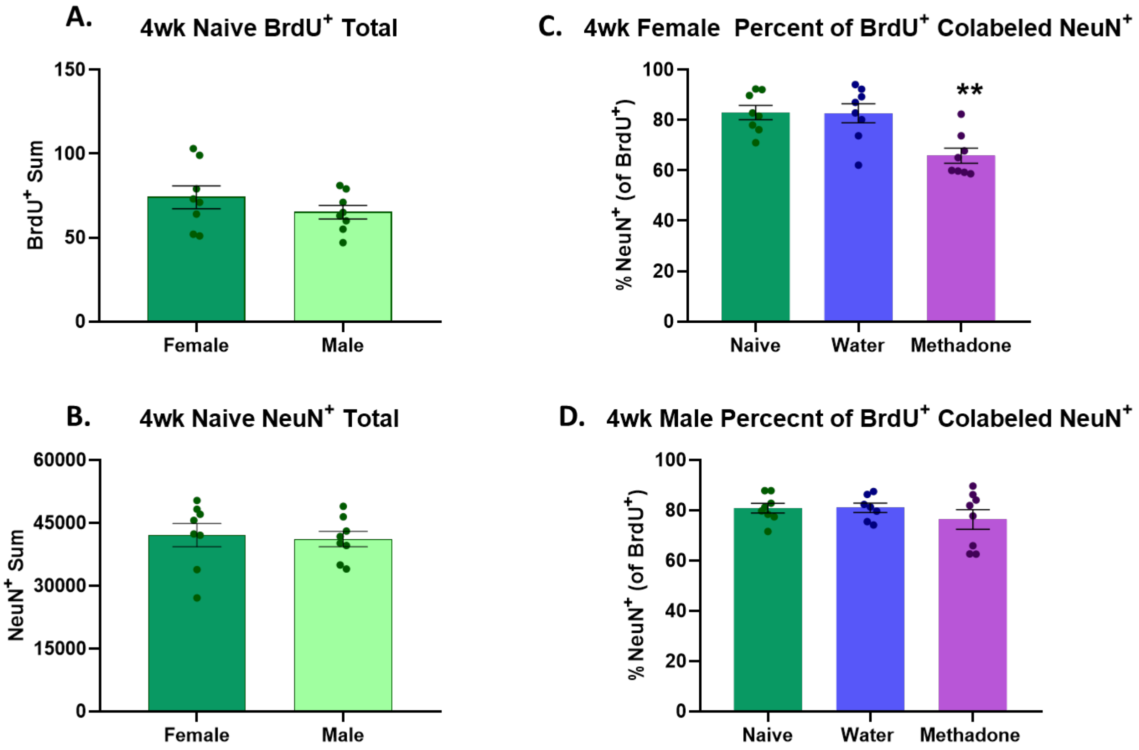


**Supplementary Figure 6.** There were no significant sex differences in the number of $BrdU^+$ cells (A) or $NeuN^+$ cells (B) in naïve controls at the 4wk timepoint. There were also no significant exposure differences in the number of $BrdU^+$ cells that were colabeled with NeuN for either sex (C-D). Data points represent individual animal counts; error bars represent SEM.
